# Why models underestimate West African tropical forest productivity

**DOI:** 10.1101/2024.03.08.584066

**Authors:** Huanyuan Zhang-Zheng, Benjamin Stocker, Eleanor Thomson, Jesús Aguirre-Gutiérrez, Xiongjie Deng, Ruijie Ding, Stephen Adu Bredu, Akwasi Duah-Gyamfi, Agne Gvozdevaite, Sam Moore, Imma Oliveras Menor, I. Colin Prentice, Yadvinder Malhi

## Abstract

Tropical forests dominate terrestrial photosynthesis, yet there are major contradictions in our understanding due to a lack of field studies, especially outside the tropical Americas. A recent field study indicated that West African forests have among the highest forests gross primary productivity (GPP) yet observed, contradicting models that rank them lower than Amazonian forests. Here, we explore possible reasons for this data-model mismatch. We found the in situ GPP measurements higher than multiple global GPP products at the studied sites in Ghana. The underestimation of GPP by models largely disappears when a standard photosynthesis model is informed by local field-measured values of (a) fractional absorbed photosynthetic radiation (fAPAR), and (b) photosynthetic traits. Satellites systematically underestimate fAPAR in the tropics due to cloud contamination issues. The study highlights the potential widespread underestimation of tropical forests GPP and carbon cycling and hints at the ways forward for model and input data improvement.

**Related manuscript:** The recent field study mentioned above is a manuscript currently accepted by *Nature Communications* (manuscript id NCOMMS-23-37419), which is available as a preprint https://www.researchsquare.com/article/rs-3136892/v1

**Codes and data availability:** All data and codes underlying the study are currently shared via [Github (link here)](https://github.com/Hzhang-ouce/Why-models-underestimate-tropical-forest-productivity) which will be made available through Zenodo upon acceptance.

## 1 Introduction

Carbon exchanges between terrestrial ecosystems, especially tropical forests, and the atmosphere are a major element of the global carbon cycle. As the world’s most productive terrestrial ecosystems ^1^, tropical forests have been estimated to account for around 44% of global forest biomass and 43% of global gross primary production (GPP) ^2,3^. Nonetheless, confidence in estimates of tropical forest productivity remains low ^4,5^. There is still a large variation among models regarding the magnitude and spatial pattern of tropical GPP ^6–8^. Multiple global-scale forest GPP studies have indicated that the largest uncertainty applies to tropical forests ^9–12^. This is likely an inevitable consequence of the paucity of carbon cycling data in the tropics relative to temperate regions ^13–16^. For instance, many previous studies found particularly large GPP data-model discrepancy for tropical forests between flux towers and models (including satellite GPP products) ^16–18^, but investigation into the causes of discrepancy was not attempted due to the lack of *in situ* auxiliary measurements of plant traits and forest characteristics ^19^. The large unresolved data-model discrepancy suggests fundamental challenges in our understanding of tropical forest productivity and its geography.

The uncertainty in estimates of tropical forest productivity is particularly large in West Africa^20^. The first field quantification of GPP in African forests (by measuring each component of GPP in three study sites, termed ‘biometric’) reported possibly the highest GPP value recorded in intact forests – greatly surpassing values measured in Amazonia ^21^. However, the high GPP of the three study sites are underestimated by about 60% in both the MODIS and FLUXCOM products, two widely used global maps of tropical forest productivity ^21^. It has not yet been explained, from modellers’ perspective, why this area has surprisingly high GPP. As models are designed based on current ecological theory ^22^, such a large discrepancy of multiple sites signals a lack of physiological understanding underpinning the high productivity. Here we set out to explain the high productivity of West African forests using photosynthesis models and investigate the reasons behind the data-model discrepancy, with a view to provide information to modellers working on assessing and mapping the productivity of forests globally.

The investigation follows three objectives. **Objective (1):** We first compare the biometric GPP to multiple models and satellite-based products, to quantify the data-model discrepancy. **Objective (2):** We investigate whether the field-observed high productivity is consistent with the leaf photosynthetic traits and other field measurements that were commonly simulated by models. This investigation was made possible by extensive field measurements of environmental variables, plant traits, and carbon fluxes at the study sites ^23–26^. **Objective (3):** We attempt to account for the data-model discrepancy in the West African region by substituting model parameters with field measurements, according to several hypotheses listed below as to the cause of this discrepancy.

Most vegetation models estimate GPP using key inputs including:

- light use efficiency (LUE), a key parameter of MODIS-GPP and P-model (see Methods section for description of models involved). LUE could not be directly measured but could be derived from field-measured photosynthetic capacity ^27^ (supplementary method). P-model ^27^ and most dynamic global vegetation models (DGVMs) use photosynthetic capacity to calculate GPP ^28^;
- fraction of absorbed photosynthetically active radiation (fAPAR), a key parameter of MODIS-GPP and P-model. Most DGVMs simulate leaf area index, which is closely related to fAPAR;
- plant functional types (PFTs) classification;
- climate variables, including temperature, relative humidity and photosynthetic photon flux density (PPFD), often calculated from incoming shortwave radiation. For models these are retrieved from global climate data products.

Therefore, we hypothesize that the large data-model discrepancy for GPP could stem from inaccuracies in one or more of these key input variables: (**Hypothesis 1**) incorrect LUE, or photosynthetic capacity; (**Hypothesis 2**) incorrect fAPAR; or (**Hypothesis 3**) inappropriate assignment of plants functional types; or (**Hypothesis 4**) climate variables. Multiple experiments are designed to test each hypothesis (Figure 3).

**Figure 1.**
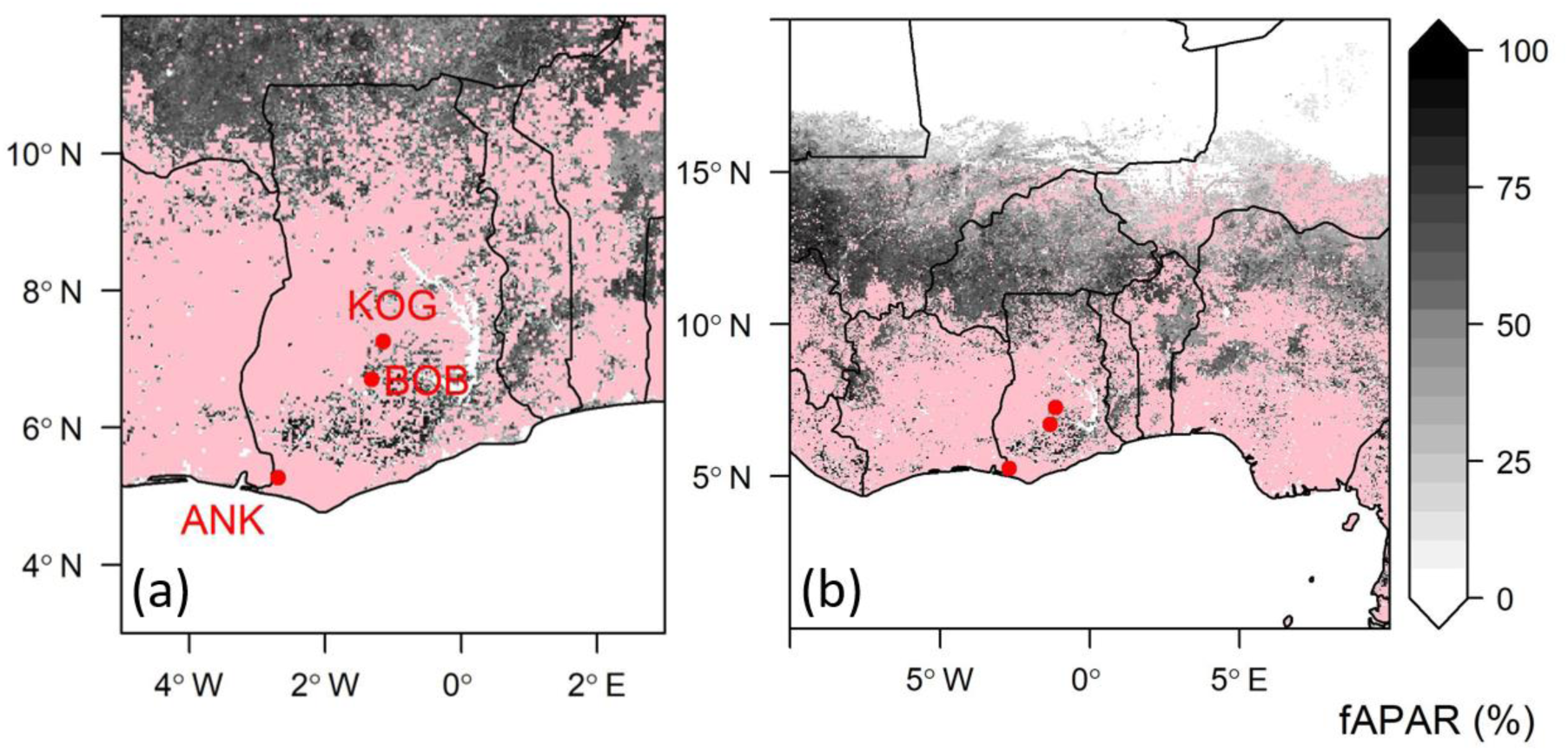
Map of the three study sites in Ghana, West-Africa. Grey scales show MODIS fAPAR, on 2016-09-07. Pinks areas are pixels contaminated by clouds, which were flagged as ‘Significant clouds were present’ in MODIS MOD15A2H product. Each red dot denotes a site. Each site contains multiple one-hectare plots (see Method). More meta-data on these sites are provided in the supplementary.

**Figure 2.**
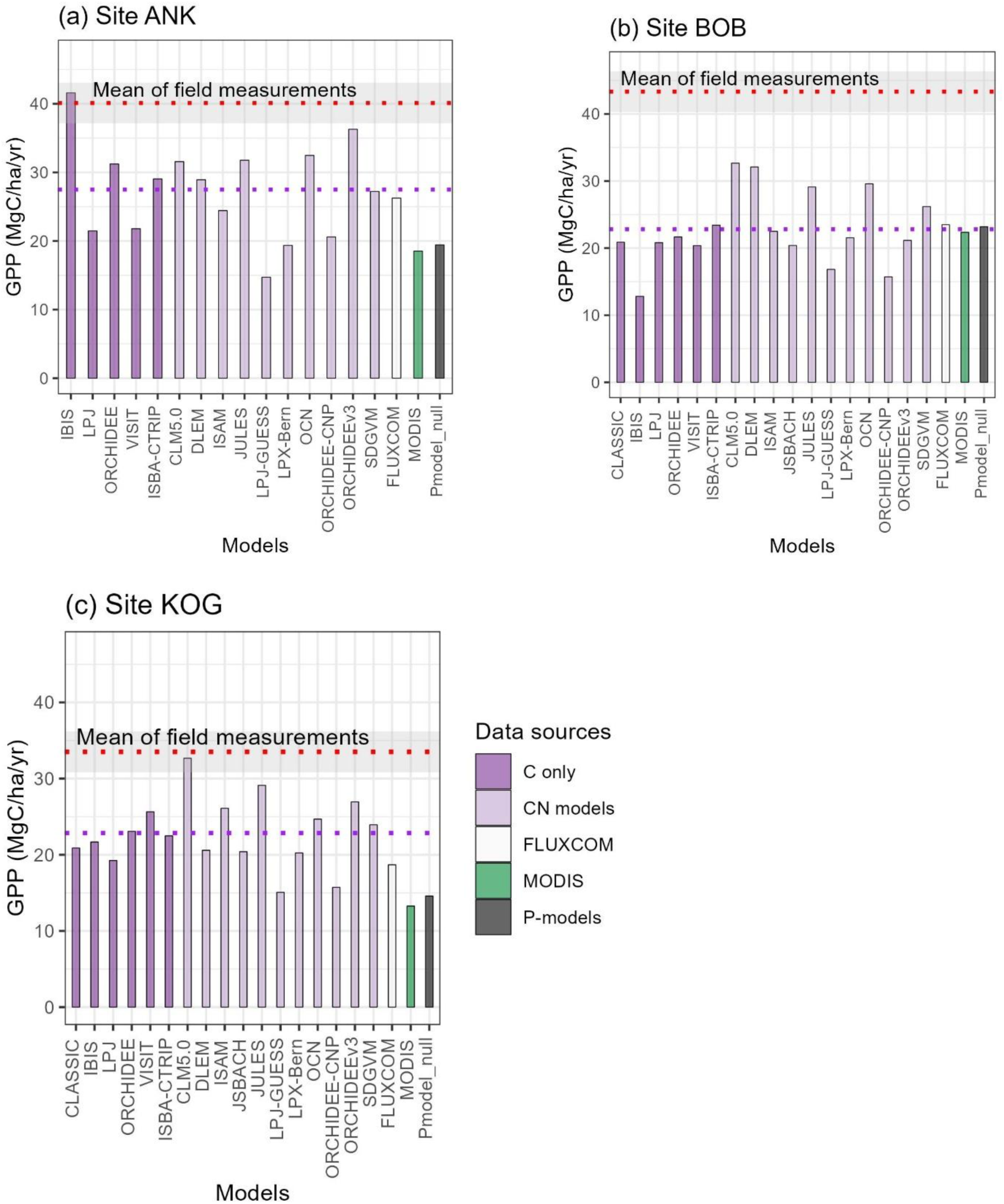
Intercomparison of gross primary productivity (GPP, MgC/ha/year) from various independent sources. The figure contains study sites (a) Ankasa, (b) Bobiri and (c) Kogaye.. The red dotted lines denote *in situ* biometric GPP, as a mean of multiple one-hectare plots (Table S3) ^21^. The grey areas denote measurement uncertainty, not standard error. The uncertainty is calculated through error propagation. Bars denote Carbon only models (dark purple), Carbon-Nitrogen coupled models (light purple), FLUXCOM (white) and MODIS (green). Purple dotted line denotes all models average.

**Figure 3.**
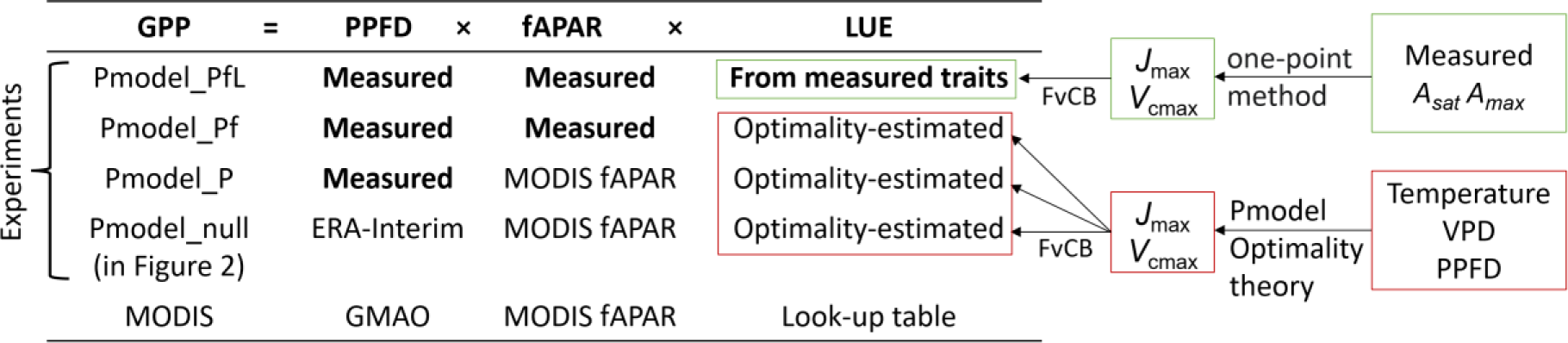
Input data for experiments in Figure 4. Input using field measurements are in bold. The schematic shows the calculation of LUE for the four experiments.

Overall, we identify key areas for model improvement and also provide a trait-based explanation for the high GPP observed in West Africa.

## 2 Methods

### 2.1 Study sites and Field measurements

The three study sites span a wet-to-dry rainfall gradient in Ghana from evergreen rainforest at site Ankasa (ANK) with a mean annual precipitation of 2050 mm, to semi-deciduous forest at site Bobiri (BOB) with 1500 mm, and a dry forest and mesic savanna matrix at site Kogyae (KOG) with 1200 mm. Each study site represents a major forest type of West African forests. See Table S3 for site environmental data.

Biometric GPP was measured in the field using the Global Ecosystem Monitoring protocol, which sums measurements of the components of NPP and autotrophic respiration to calculate GPP ^26^ and is described in detail in a previous study ^21^. Biometric GPP was originally quantified at the plot scale. There are three one-hectare plots at Ankasa, six plots at Bobiri, and five plots at Kogyae. This study was conducted at site rather than plot scale because one-hectare plots at the same site share almost identical climate variables and fall into one model grid cell. Nonetheless, plots at the same site fall into different grid cells of the MODIS product (Figure S 1). We therefore calculated, for each site, the mean biometric GPP across plots, but not standard error. The uncertainty in Figure 2 represents measurements error as calculated by error propagation from each GPP component, instead of spatial variation of GPP. See Supplementary method and ^21^ for more data and error propagation associated with the study sites.

The fraction of absorbed photosynthetically active radiation, fAPAR, was obtained using hemispherical photography, taken each month in each plot between 2012 and 2017. Photos were processed using machine learning-based software ‘*ilastik*’ ^29^ for pixel classification and CANEYE ^30^ for fAPAR calculations (Supplementary Methods).

In this study, we also need field-measured community-weighted mean light-saturated photosynthetic rate (*A*_sat_), light- and CO_2_-saturated photosynthetic rate (*A*_max_), to calculate a trait-based LUE and trait-based GPP (specific experiment explained in Section 2.4). We derived *V*_cmax_ and *J*_max_ from each field measured *A*_sat_ and *A*_max_ at mean annual air temperature using R package ‘plantecophys’ ^31,32^. We used these *V*_cmax_ and *J*_max_ in the simulation ‘Pmodel_PfL’ (see Section 2.4). However, the *J_max_* limitation equation in ‘plantecophys’ is different to that in ‘*rpmodel*’ (an R package used for experiments explained below), so we modified the *J_max_* equation in ‘*plantecophys*’ (see Supplementary Methods). Measurements of photosynthetic traits were made every three months from 2014 to 2016 to cover both wet and dry seasons. Although we measured both shade and sun leaves, we used sun leaves only in this study, consistent with common practice in field studies of photosynthetic traits ^33^. Besides, we used above-canopy PPFD as model input and only sun leaves acclimate to this level of PPFD; shade leaves acclimate to darker environments and have consistently lower *A*_sat_ than sun leaves^34^.

### 2.2 Compare biometric GPP to models’ GPP (Figure 2)

For **Objective (1)**, we compared biometric GPP with values estimated by (a) the TRENDY ensemble of dynamic global vegetation models (DGVMs) ^35^; (b) two data-driven models, FLUXCOM ^36^, and MODIS ^37^, and (c) an optimality-based model (P-model v1.0) ^27^. The above choices are the most extensively used representatives from each GPP method category, according to a thorough review of many GPP products by Zhang & Ye ^16^.

The TRENDY ensemble is a collection of 15 DGVMs that calculate functional aspects of vegetation, including fAPAR or leaf area index (LAI), metrics that determine light absorption from environmental variables, without using any remotely sensed data as input. Most of these models calculate leaf-level photosynthesis via the Farquhar-von Caemmerer-Berry (FvCB) photosynthesis model ^38^, which requires specification of several photosynthetic traits: the maximum carboxylation rate (*V*_cmax_), the maximum electron transport rate (*J*_max_), and parameters of one or other of the semi-empirical schemes that are commonly used to estimate stomatal conductance (*g*_s_) ^28^. Leaf-level photosynthesis is scaled up to the canopy, and thus to GPP, by methods that vary in complexity, but all depend on the modelled LAI or fAPAR.

FLUXCOM and MODIS, by contrast, are data-driven models that do not depend on the FvCB model. FLUXCOM GPP is a machine learning application that uses eddy-covariance estimates of GPP from worldwide flux towers as the key input, combined with environmental covariates that include satellite-derived fAPAR, and shortwave radiation (closely related to PPFD). FLUXCOM used MODIS fAPAR, and shortwave radiation prepared by Japan Aerospace eXploration Agency (JAXA) using Terra MODIS data.

MODIS GPP ^37^ is a light use efficiency model ^39,40^ that calculates GPP as:

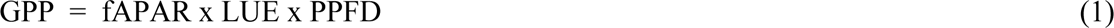

where PPFD is sourced from Global Modeling and Assimilation Office (GMAO). The MODIS GPP algorithm calculates LUE as a prescribed (per biome) maximum light use efficiency, multiplied by reduction factors that are defined *a priori* as biome-specific functions of temperature and vapour pressure deficit.

In this analysis we also employ Pmodel, R package ‘rpmodel’ ^27^. The Pmodel, uniquely, is a LUE model (also using Equation 1) but it calculates LUE based on the FvCB model, using optimality principles ^41^ to determine the spatial and temporal variation in *V*_cmax_, *J*_max_ and the ratio (c_i_/c_a_) of leaf-internal to ambient CO_2_ ^27^. The c_i_/c_a_ ratio results from the combined effects of photosynthetic rate and stomatal conductance, which are co-regulated by plants. Leaf-level photosynthesis is scaled up to the canopy with the help of the big-leaf approximation ^42^ and driven by satellite fAPAR data. The Pmodel thus combines the mechanistic basis of photosynthesis as represented in DGVMs with the simplicity of LUE models. The Pmodel also dispenses with the need to consider plant functional type (PFT) or biome distinctions (apart from the difference between C_3_ and C_4_ plants); the differences in photosynthetic traits among C_3_ PFTs are implicitly predicted as a consequence of their habitat preferences alone. The validity of these predictions has been supported by global-scale comparisons ^43–45^.

### 2.3 Consistency between the in situ biometric GPP, and other in situ measurements

Before investigating the cause of the GPP data-model discrepancy **(Objective 3)**, it is necessary to first investigate whether biometric GPP can be reproduced by the FvCB model (within Pmodel) fully informed by field-measured inputs – PPFD, fAPAR and photosynthetic capacities **(Objective 2)**. Here we fed Pmodel with *V*_cmax_ *J*_max_ derived from field measured *A*_sat_ *A*_max_ (See equations and codes in supplementary method). Note that the ‘big leaf assumption’ is implied for all Pmodel simulation in this study. Pmodel calculates a LUE from *J*_max_ before calculating GPP. The above GPP experiment was called ‘Pmodel_PfL’. A match would indicate consistency between the field-measured canopy properties and the independent field biometric GPP, strengthening confidence in the field estimates.

If such a match is found between Pmodel_PfL GPP and biometric GPP, we can further investigate the cause of GPP data-model discrepancy by designing more experiments using Pmodel with different inputs (see 2.4 and Figure 3). The difference between Pmodel_PfL and biometric GPP is labelled as the **’unresolved discrepancy’** in Figure 4.

**Figure 4.**
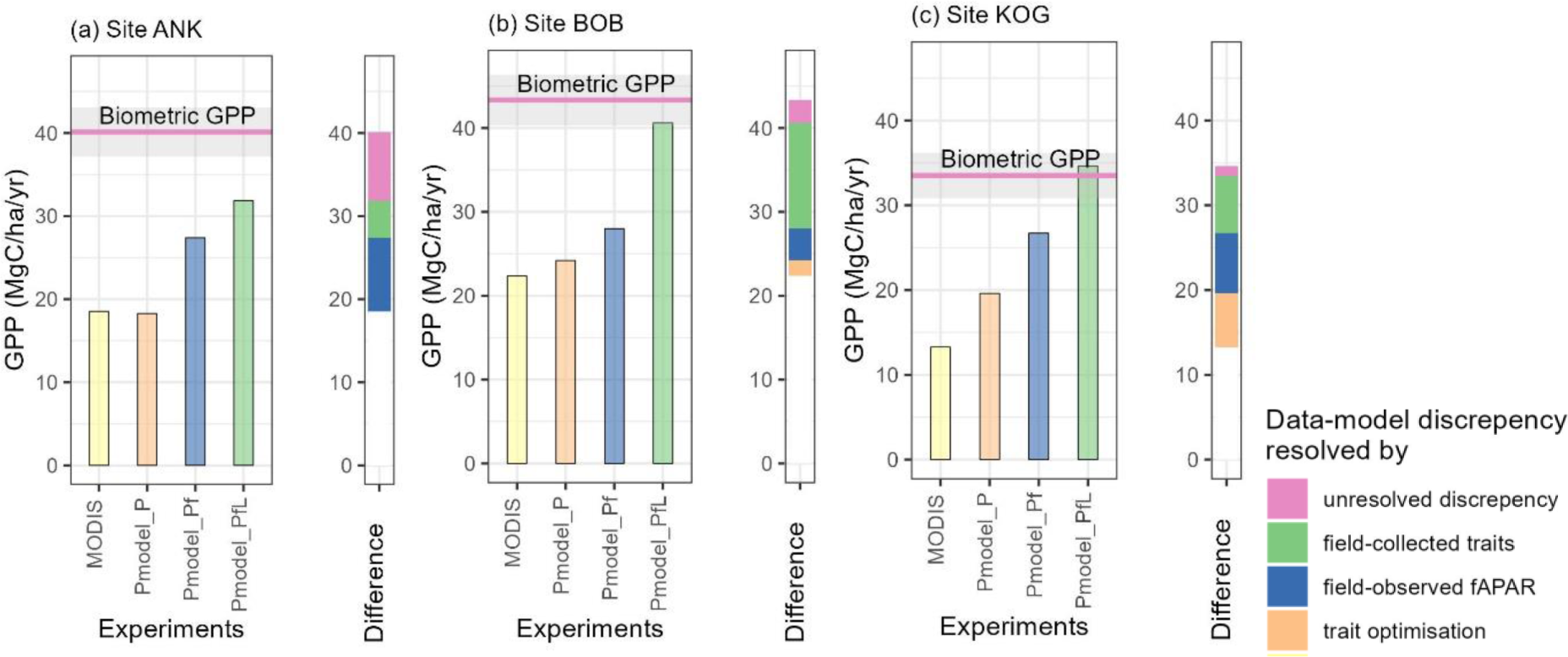
Partitioning of GPP data-model discrepancy, by comparing different experiments. Results are shown for study sites (a) Ankasa, (b) Bobiri and (c) Kogaye. The left panel displays field-based biometric GPP (pink line with grey zone showing uncertainty) and multiple GPP experiments (bars). Pmodel_PfL, Pmodel_Pf and Pmodel_P are GPP experiments which are all simulated using P-model but with different inputs (see Figure 3 and Methods). Both P-model and MODIS GPP were calculated from equation (1), enabling direct comparison. The right panel, a ‘Diff’ bar, illustrates the difference in GPP between experiments, which represents the sources of GPP data-model discrepancy. For example, the GPP difference between Pmodel_Pf and Pmodel_P experiments (Figrue 3) is caused solely by the difference in input variables - fAPAR. Therefore, the difference between them (blue) is GPP data-model discrepancy caused by fAPAR. The difference (orange) between Pmodel_P and MODIS is GPP data-model discrepancy caused mainly by the difference between optimality-based LUE and MODIS LUE. The difference (green) between Pmodel_PfL and Pmodel_Pf is GPP data-model discrepancy caused by the difference between measured LUE and optimality-based LUE.

### 2.4 Cause of GPP data-model discrepancy

For **Objective (3)**, following equation 1 (also see Introduction), we hypothesize that the large data-model discrepancy for GPP could stem from: (**Hypothesis 1**) incorrect LUE, or photosynthetic capacity; (**Hypothesis 2**) incorrect fAPAR; or (**Hypothesis 3**) inappropriate assignment of plants functional types; or (**Hypothesis 4**) climate variables. This hypotheses testing is illustrated in Figure 3.

To test Hypothesis 1 (incorrect LUE explains the mismatch), we used Pmodel to predict photosynthetic capacity (Vcmax, Jmax) and LUE using optimality theory based on climate variables alone. Pmodel then calculates a ‘Pmodel_Pf’ GPP using the above optimality-based predictions, combined with field-measured PPFD and fAPAR. As the only difference between ‘Pmodel_Pf’ and ‘Pmodel_PfL’ GPP is in light use efficiency (derived either from optimality or from *in situ* measurements), we label the difference in GPP as **’data-model discrepancy resolved by field traits’**

Next, the model calculates a ‘Pmodel_P’ GPP also using measured PPFD and the above optimality-theory predicted LUE, but with MODIS-derived fAPAR rather than *in situ* measured fAPAR as input. This ‘Pmodel_P’ GPP is compared to MODIS GPP. We note that a small proportion of the difference between MODIS GPP and Pmodel_P might originate from different PPFD inputs, because Pmodel_P uses field PPFD whereas MODIS GPP uses PPFD from NASA GMAO (Figure 3) ^37^; however, the differences are expected to be slight (see Figure S 3). The primary difference between ‘Pmodel_P’ GPP and MODIS GPP is in LUE (predicted by optimality theory versus from MODIS lookup table), hence the difference is labelled as **‘data-model discrepancy resolved by optimality’**. The secondary difference between ‘Pmodel_P’ GPP and MODIS GPP is in PPFD. The secondary difference is neglected because the PPFD used by MODIS is not publicly available. Also, we showed that satellite PPFD is similar to measured PPFD (Figure S 3).

To test Hypothesis 2 (incorrect fAPAR explains the mismatch), we compared ‘Pmodel_P’ to ‘Pmodel_Pf’. The only difference between these two is in fAPAR (*in situ* measurements versus MODIS). We referred to the difference as **’data-model discrepancy resolved by fAPAR’.**

Lastly, ‘Pmodel_null’ GPP is calculated using satellite PPFD, in which case Pmodel gets no information from field measurements, which is a fair comparison to other models (Figure 2).

Equivalently to comparing GPP, one could directly compare *V*_cmax_, *J*_max_, fAPAR and LUE from multiple sources (some presented in Figure S 2 and Figure S 3). However, LUE and GMAO PPFD used in MOD17 GPP are not publicly available and were difficult to reproduce, so we did not include them. Since the data-model discrepancies in these variables all cascade into GPP, we focus on visualizing the discrepancies in GPP in Figure 4.

Hypothesis 3 (confusion of land cover classification) is based on the fact that the GPP estimated from *in situ* measurements, which is the mean of data from several one-hectare plots, differs in scale from the GPP estimated by the TRENDY models, FLUXCOM and MODIS (as visualized in figure S1). West African forests are extremely fragmented, and some forest patches are smaller than the grid cell size of the models. For example, one of the study forests, Bobiri (BOB), is a forest fragment measuring only about 7 x15 km (Figure S 1), surrounded by cocoa farmland. To illustrate this, we drew a map of MODIS GPP (with a resolution of 500m) over the Bobiri site, covering an area similar to a single 0.5° x 0.5° grid cell of the TRENDY models. Since each grid cell of the TRENDY models is a composite of multiple PFTs (often represented by different fractions in different models) and models report GPP per PFT, we also show the modelled PFT composition of the grid cells and compare ‘forest-only’ GPP to the grid cell average (Figure S 4).

### 2.5 Data-model comparison for climate variables and c_i_/c_a_

For **Objective 3 Hypothesis 4**, the following analysis was conducted. The ratio c_i_/c_a_ was estimated from leaf δ^13^C measurements, using the method described in ^46^. This isotope-derived c_i_/c_a_ was compared to Pmodel-predicted c_i_/c_a_ (Figure S 3). The Pmodel requires several climate variables as input, and this can be a source of uncertainty in modelled GPP. To address this, we compared the temperature and relative humidity from local weather stations to products commonly used by vegetation models. We selected relative humidity and temperature provided by ERA-interim, from which we also calculated optimality-estimated c_i_/c_a_ via ‘*rpmodel*’. We derived PPFD from shortwave radiation retrieved from the ERA5-Land climate reanalysis product, following ^43^.

Previous data-model comparisons of climate variables for tropical forest sites (Burton, Rifai and Malhi, 2018) found ERA-interim to be the best product for tropical Africa. As we found very small data-model discrepancy in climate variables (see Results), we used ERA-interim temperature, vapour pressure and optimality-estimated c_i_/c_a_ for all GPP experiments in this study to maintain consistency and ensure a fair comparison between P-model and other models, while the choice of PPFD dataset was specific to the experiment (see simulation schematic in Figure 3).

### 2.6 MODIS fAPAR and GPP

Using Google Earth Engine, we retrieved MODIS GPP from the collection MOD17A2H and retrieved MODIS fAPAR from MOD15A2H over 2001 to 2020. We extracted GPP and fAPAR of the 14 plots using their coordinates and calculated annual (Figure 2 and Figure 4) and monthly mean values (Figure 5) per site. Plots in the same site fall into one grid cell of TRENDY model (approximately 50 x 50 km, or 0.5° x 0.5°), but into different grid cells of MODIS due to higher resolution (500m). The scale difference is illustrated in Figure S 1. To remove cloud-contaminated GPP and fAPAR, we selected only data flagged as ‘Significant clouds NOT present (clear)’ or ‘Mixed cloud present on pixel’. Since MODIS GPP and fAPAR share the same data flag, the above approach ensures that the fAPAR used in Pmodel_P and Pmodel_null is identical to that used in MODIS GPP (Figure 4), ensuring fair comparison between them. The issue of cloud contamination is visualized in Figure 1 and Figure 5.

**Figure 5.**
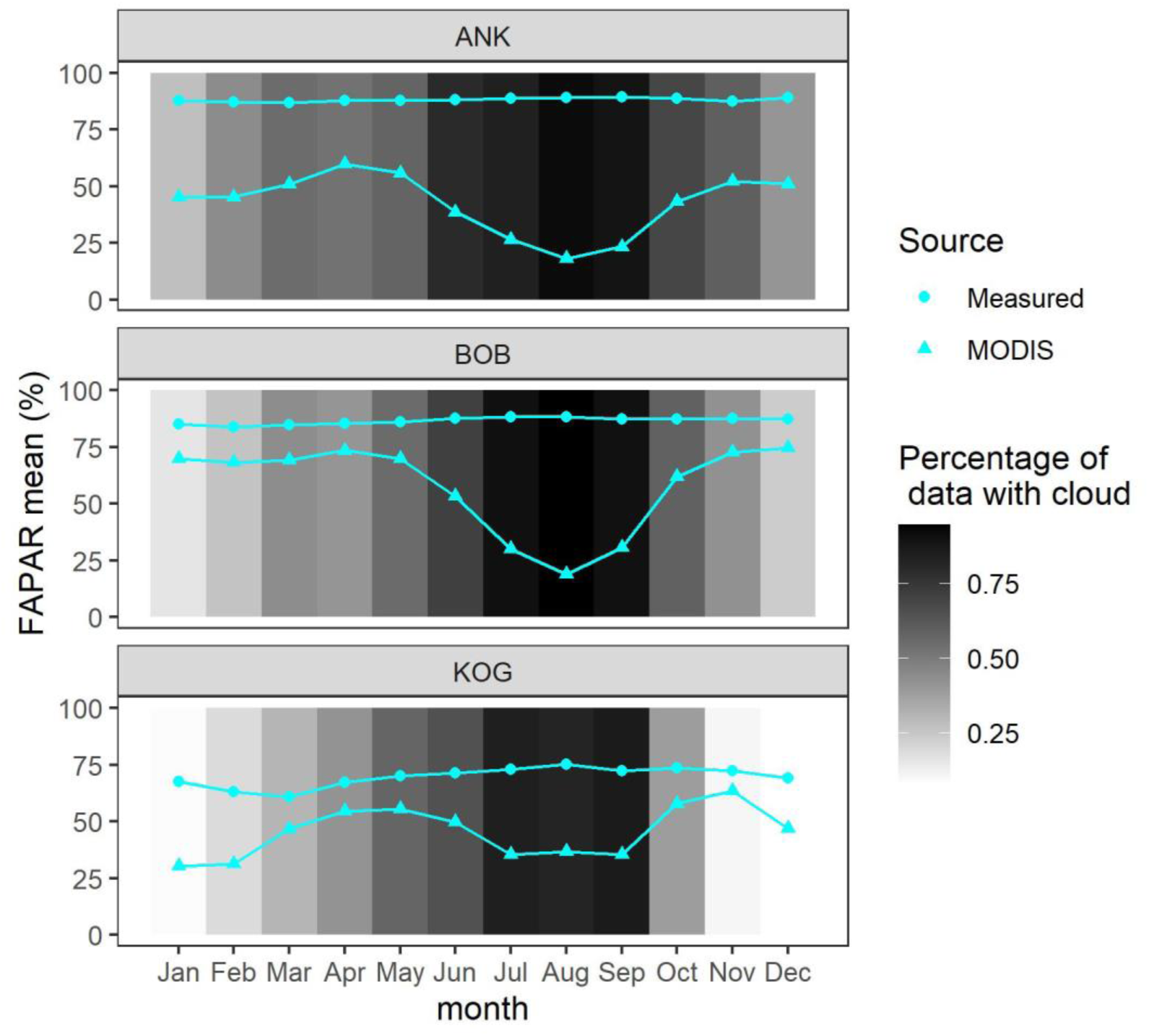
The seasonal variation of the fraction of absorbed photosynthetically active radiation (fAPAR). The figure presented data retrieved from MODIS MOD15A2Hv006 (triangle) and *in situ* measured using hemispherical photography (dot), at the three study sites (ANK, BOB and KOG), as a monthly mean across 2001 to 2020. Note that fAPAR in the above figure was displayed without cloud contamination filtering (nor gap-filling), but MODIS fAPAR used in Figure 3 is filtered. Grey scale shows the percentage of records contaminated by clouds.

### 2.7 FLUXCOM and TRENDY GPP

There was a flux tower at ANK (GH-Ank), but not at BOB or KOG. Therefore, we compared biometric GPP at the three sites with FLUXCOM. We chose the ‘RS_METEO’ version of FLUXCOM because the magnitude of GPP in this version does not involve uncertainty from MODIS fAPAR, which makes the comparison between FLUXCOM and MODIS GPP more independent. For TRENDY, we analysed the model outputs in version 9 ^35^ under the S2 protocol, in which climate and CO_2_ change while land use is kept constant. Models are classified into nitrogen-carbon coupled models and carbon only models for readers convenience. Note that models still include ‘C4 crop land’ as one of the plant functional types (PFTs), but the cover of ‘C4 crop land’ is kept constant. We retrieved total GPP (sum of all PFTs) of the grid cell (variable ‘gpp’). We also used variables ‘gpppft’ and ‘landcoverfrac’ to calculate a forest-only GPP, which is the GPP that the model would have estimated if the whole pixel were ‘forests’, including evergreen, deciduous and any other types of forests. For both TRENDY and FLUXCOM, we extracted the grid cells where the three study sites are located, as an average from 2009 to 2019.

## 3 Results

### 3.1 Intercomparison of GPP estimates

Considerable discrepancies in GPP are found among DGVMs, FLUXCOM, MODIS, Pmodel, and biometric measurements for all sites, with a pattern that is consistent from site to site (Figure 2). At each site, biometric measured GPP exceeds GPP estimated by other methods. Overall, the DGVMs models’ average GPP is higher than FLUXCOM, which in turn is higher than MODIS GPP. At BOB, the DGVM average, FLUXCOM, and MODIS report almost equal GPP – but still less than the measured GPP. There are substantial disagreements between different DGVMs; for example, at ANK, the DGVMs simulated GPP ranged from 15 to 42 (biometric GPP at 40) MgC/ha/year. The largest data-model discrepancy between DGVMs and biometric GPP is seen at BOB (22.8 versus 43.3 MgC/ha/year). The discrepancy between biometric and MODIS GPP is about 20 MgC/ha/year at all sites.

### 3.2 The match between trait-based GPP and biometric GPP

Using field observed PPFD, fAPAR, and photosynthetic traits as input, Pmodel could estimate a GPP (i.e., experiment Pmodel_PfL) slightly lower but still within the uncertainty range of field biometric GPP at BOB and KOG sites. At the ANK site, Pmodel_PfL greatly reduces the data-model discrepancy but remains short of biometric GPP (Figure 4). Nonetheless, the slight mismatch at ANK does not undermine further investigation at ANK because there are more dominant factors contributing to the data-model discrepancy, explained below.

### 3.3 Measured and modelled photosynthetic trait

Photosynthetic capacity estimated by the Pmodel is lower, although close to field measured photosynthetic capacity (Figure S 2) implying underestimated LUE. As a consequence, Pmodel_Pf is lower than Pmodel_PfL consistently at all sites (Figure 4).

GPP is substantially underestimated when *in situ* fAPAR is replaced with MODIS fAPAR, resulting in a considerable difference between Pmodel_Pf GPP and Pmodel_P at any site. Especially at ANK, fAPAR is found to be the largest contributor to GPP data-model discrepancy. A further investigation into MODIS fAPAR (Figure 5) shows that during the rainy season, MODIS fAPAR decreases along with increasing cloud cover (up to 90% of pixels), whereas field fAPAR peaks (due to a denser canopy) during the rainy season. In short, MODIS fAPAR shows an opposite trend to field fAPAR due to cloudiness. Filtering fAPAR values according to cloudiness or gap-filling of bad data points would remove most of the values from the rainy season, leading to an underestimated mean annual fAPAR. The above issue affects a major proportion of the African tropical forest, and likely many areas of the tropics (Figure 1).

At the ANK site, MODIS and Pmodel_P GPP are almost identical, but at BOB and KOG MODIS is much lower than Pmodel_P GPP. The difference could originate from PPFD (Figure S 3) or because the optimality-based LUE is higher than MODIS LUE (from the look-up table). The PPFD and LUE used by MODIS GPP are not publicly available and therefore a definitive conclusion cannot be given. Considering that Pmodel_null (using ERA5 PPFD) is also higher than MODIS GPP at any site (Figure 2), it is more probable that optimality theory tends to estimate an LUE higher than LUE in MODIS GPP.

In short, Hypotheses 1 and 2 are accepted.

### 3.4 Scaling and plant functional type

At BOB the entire forest fragment is smaller than a single grid cell in a DGVM. MODIS GPP estimates that the surrounding area (a composite of savanna and cocoa farms) has only half the productivity of the forests. Therefore, the mean of the whole map (19.6 MgC/ha/year) appears to be smaller than that of the forests (23 MgC/ha/year) (Figure S 1). However, field studies of net primary production (NPP) indicate that cocoa farms are as productive as adjacent forests ^47^.

For all of the study sites, most TRENDY models treat the grid cell as a composite of ‘forest’ or ‘C4 grass’, although the fraction of forests varies considerably among models (Table S1). Having a proportion of C4 grass is true for BOB (Figure S 1) and KOG, but ANK is entirely forest. Furthermore, most TRENDY models correctly estimate that ‘C4 grass/crop’ is as productive as forests (different from MODIS) and hence, forest-only GPP is similar to the mean of the whole pixel (Figure S 4). The above lead to the conclusion that the GPP data-model discrepancy between TRENDY and biometric GPP cannot be explained by scaling or PFT issues (rejecting hypothesis 3). The data-model discrepancy between MODIS and biometric GPP at the 0.5° scale, however, is compounded by MODIS’ estimation of low productivity in croplands (Figure S 4).

### 3.5 Data-model consistency in climate variables

The data-model differences we found for temperature, relative humidity, PPFD and c_i_/c_a_ were very small and not likely to contribute significantly to the GPP data-model discrepancy (Figure S 3), thus rejecting hypothesis 4.

To sum up, the data-model discrepancy for GPP at the study sites mainly stems from: incorrect LUE, or photosynthetic capacity (Hypothesis 1 accepted); and incorrect fAPAR (Hypothesis 2 accepted); **not** inappropriate assignment of plants functional types (Hypothesis 3 rejected); **nor** climate variables (Hypothesis 4 rejected).

## 4 Discussion

### 4.1 Main sources of GPP data-model discrepancy

We not only demonstrate a significant data-model discrepancy in West African forest GPP but also reveal the main sources of this discrepancy. At the Ankasa wet rainforest site, we found that underestimation of fAPAR, likely because of cloud contamination, is the major contributor to GPP data-model discrepancy. At the Bobiri (semideciduous) and Kogyae (dry forest) sites, too low values of photosynthetic traits (leading to a bias in light use efficiency) are the primary source of GPP data-model discrepancy, which can be partly explained using optimality theory predictions. Additionally, fAPAR accounts for a large proportion of GPP data-model discrepancy at KOG but not at BOB. Through this analysis we are able to fully account for the data-model discrepancy at KOG and BOB, but leave a modest, though greatly reduced, unresolved data-model discrepancy still persisting at ANK, which may be caused by photosynthesis model assumption (e.g. the big leaf assumption) or possible bias in field measurements ^48^.

### 4.2 Why do models underestimate GPP in West African tropical forests?

Focusing on three sites (comprising 14 one-hectare plots) in Ghana, we have utilized the first *in situ* quantification of African forest GPP and conducted a systematic data-model comparison. Despite the uncertainty associated with biometric GPP measurements, it is more likely that FLUXCOM, MODIS, TRENDY models, and Pmodel_null underestimate West African forest productivity than that the biometric GPP measurements overestimate it. This is because (1) the biometric GPP is an average across multiple plots spanning several years, and its high productivity is supported by another field study using similar biometric method of GPP quantification ^47^; (2) from a photosynthesis traits perspective, West African seasonally dry forests are characterised by a high CO_2_ assimilation rate or high photosynthetic capacity, in comparison to wet-evergreen tropical forests studied in other continents, predominately in Amazonia ^25,49,50^; P-model could simulate a GPP close to biometric GPP if informed by field-measured traits (Figure 4), which signals a broad consistency between the high GPP and observed photosynthetic traits. (3) the cause of the models’ underestimation can be well explained by errors in modelled LUE and satellite-derived fAPAR (Figure 4).

Some degree of model underestimation of GPP may even be a pan-tropical feature. Intercomparison of GPP and NPP in previous studies ^51,52^ has also revealed data-model discrepancies for Amazonian lowland forests. In this study, we found that the data-model discrepancy was due to fAPAR and photosynthetic capacity. The cloudiness issue, leading to low satellite-based fAPAR, has been described as a pan-tropical phenomenon for many remotely sensed products ^53–56^. Bias in MODIS fAPAR (or LAI) will inevitably cascade into models or analyses that use MODIS fAPAR as a predictor, including FLUXCOM. Similar to our findings, Wang *et al* indicated that fAPAR (and not meteorological data) is the main cause of MODIS GPP underestimation in both African and Amazonian forests ^57^. Many West and Central African forests have high persistent cloud cover compared to Amazonian forests and therefore may be disproportionally affected by cloud contamination.

The LUE (or photosynthetic capacity) discrepancy is strong and consistent with previous literature, since a previous study found higher *A*_sat_ and *A*_max_ values in West African species than in Amazonian ones ^50^. Higher *A*_sat_ and *A*_max_ are found in BOB and KOG (drier sites), but not ANK (the wet evergreen forest on poor soils more similar to much of lowland Amazonia) ^21,25^, making LUE the dominating source of GPP bias in BOB and KOG but less dominant in ANK. The high photosynthetic capacity in semi-arid forest or savanna is consistent with other field studies in West Africa ^49^. *V*_cmax_ inferred from remotely sensed leaf chlorophyll data and *V*_cmax_ predicted by the P model both show exceptionally high values in West Africa and parts of India – substantially higher than in Amazonia or SE Asia ^58,59^. These seasonal forests on more fertile soils may have photosynthesis rates optimized to high light, temperature and VPD (also see field study ^60^). This substantial spatial variability of *V*_cmax_ has not been incorporated in most TRENDY models ^28,61^ which could lead to underestimation of GPP at such regions.

Rogers *et al* recalculated plant functional type specific *V*_cmax_ for numerous models (Table S2) ^62^, providing further insights into model parameterization, but the *V*_cmax_ recalculated by that study may not correspond to the *V*_cmax_ behind TRENDY v9 simulation (Figure 1 and Table S1) because the model version or simulation scenario may be different (*V*_cmax_ behind TRENDY simulation is not directly retrievable). As shown in Table S2, two TRENDY models (OCN and ORCHIDEE) indeed underestimate *V*_cmax_. IBIS substantially overestimates *V*_cmax_ which results in its high GPP at site ANK. However, JULES, CLASSIC, CLM5.0 and JSBACH have similar or higher *V*_cmax_ but still underestimate GPP, which implies that there are other factors contributing to the GPP underestimation. Moreover, the substantial inter-model disagreement shown in Table S2 is alarming, with *V*_cmax_ for evergreen forest vary from 18 to 163 μmol m^−2^ s^−1^ while field measurements at our study sites are around 30 μmol m^−2^ s^−1^ ^63^. Model parameterizations of *V*_cmax_ for C4 grass also vary substantially and appear to be lower or higher than forests depending on models, which may affect some models that have C4 grass as main plant functional types at the study sites, for example CLM5.0. To sum up, the investigation into the cause of DGVMs underestimating GPP is not straightforward. The parameterization of *V*_cmax_ is one of the reasons for some models but there are likely other contributing factors that require future studies.

### 4.3 The importance of field evidence in productivity estimation

The availability of comprehensive field measurements allows us to trace and quantify the sources of data-model discrepancy in the GPP of West African forests. We find that fAPAR and LUE (or photosynthetic capacities) are the dominant sources of data-model discrepancy, rather than model structure or climate variables. These findings reflect the lack of field measurements of West African forest photosynthetic traits and leaf area index (or fAPAR), while the environmental variables of the study regions are better represented. FLUXCOM also struggles with the number of field stations in the study region. There is only one flux tower for West African forests (site GH-ank at the Ankasa site), which reports three years of GPP varying dramatically ^57^. The tower reported patchy data for 2011, 2012 and 2014 with an estimated mean annual GPP at 36.06, 22.02 and 26.1 MgC/ha/year respectively (source: FLUXNET2015, variable GPP_NT_CUT_REF) ^64^. The mean GPP across three years appears to be lower than the biometric GPP, which may be associated with raw data quality ^65^, technical issues ^66,67^, and challenging subcanopy CO_2_ storage estimation ^68,69^. During FLUXCOM extrapolation, the machine learning algorithm would not receive information about BOB’s higher photosynthetic capacity and would simply predict BOB less productive than ANK (opposite the observed GPP pattern) due to the lower precipitation received at BOB (Table S3). Beyond pointing out the sources of data-model discrepancy, this study also highlights that such issues could be fixed if models were better informed with field-measured fAPAR and LUE derived from measured traits, or more generally if better maps of canopy traits are applied ^70^. The data-model discrepancy could also be partially (although not completely) resolved with GLASS fAPAR products ^57^ or using predicted LUE by optimality theory (Figure 4), which highlight the importance of using ‘trait-based approaches’ photosynthesis parameters instead of PFTs prescription ^28,71^. Given that GLASS, optimality-based LUE, and FLUXCOM all require validation with field evidence, this study suggests that providing models with ample field evidence and ensuring strong fidelity to field measurements is critical in improving current carbon cycle simulations ^72^.

### 4.4 Implication for carbon cycle modelling of West African forests

Researchers studying tropical forest functioning hould exercise caution when using satellite-based products subject to cloud contamination, as they are strongly compromised during the rainy season in this region due to high cloudiness. Additionally, West African forests are highly fragmented ^73^ (Figure S 1), so modellers should bear in mind that resample satellite products to a coarser resolution (e.g., from 500m to 50km) would mix forests with farmland which could be similar in productivity but differ in other aspects of carbon cycling (Figure S 4) ^47^.

The study has shown that the P-model has advanced prediction of c_i_/c_a_, *V*_cmax_ and *J*_max_ – essential parameters for the FvCB model. The data also support the coordination of the Rubiscio- and electron transport-limited photosynthetic rates, *A*_C_ and *A*_J_, which is one of the optimality principles underlying the P-model (Figure S 2). However, the reasons why almost all TRENDY models underestimate West African GPP have not been fully elucidated. Because GPP (Figure 2) and *V*_cmax_ (Table S 2) vary considerably, it is like that the causes of data-model discrepancy vary among models. While much criticism of TRENDY models has centred on issues such as the representation of nutrient limitation ^74^ or inaccurate characterizations of plant functional types ^28^, our study suggests that neither of these factors is likely to be the main reason for the underestimation of GPP at the study sites. More transparency in mode’ input data (e.g. GMAO PPFD) and parameterization (e.g. *V*_cmax_) is necessary for further investigation. Although it is challenging to unbox each model and investigate their *V*_cmax_, *J*_max_ or LUE, our analysis suggests that the underestimation of GPP is associated with photosynthetic capacities, especially at site BOB and KOG (Figure 2). Allowing *V*_cmax_, *J*_max_ or LUE to acclimate to brighter and drier environments should improve simulation of GPP at semi-arid forests and savanna ^75^.

### 4.5 Conclusion

In conclusion, the study not only reveals underestimation of West African forest productivity but also explains why this underestimation occurs. The unique data-model comparison approach proposed in this study may also have wider potential as it successfully shows (1) consistency between field measured photosynthetic traits and biometric GPP; (2) the source of data-model discrepancy by designing multiple experiments with a minimal photosynthesis model (Pmodel). The study also demonstrates that to gain insight, thorough field measurements of forest plots are valuable and application of simple models (that can be easily understood and tuned) are key for ecological hypothesis testing. As this study is limited in terms of spatial cover, we encourage future research in this region, in particular drier African ecosystems where C4 grasses are abundant with more LUE but less cloudiness (Figure 1) ^76^. We also acknowledge that models are intended for global simulation and thus our site scales study does not serve as models benchmarking but approaches to improve model simulation. Nonetheless, given the broad consistency in our results displayed across the study sites, we expect that models’ carbon cycle simulation in West African region could be substantially improved by simulating a higher GPP across West African tropical forests. It is possible, indeed likely, that such model-data discrepancies are a more general feature of tropical forests. This requires further detailed comparison between biometric field measurements and model predictions, with the approach outlined here offering a valuable approach for such a pantropical analysis.

## 5 Acknowledgement

We acknowledge the TRENDY project and participating modelling groups for the DGVM data. Y.M. is supported by the Jackson Foundation and the Leverhulme Trust. H.Z. received the Henfrey Scholarship (by St Catherine’s College, Oxford) and the Tang Scholarship (by the China-Oxford Scholarship Fund). This work is a product of the Global Ecosystems Monitoring (GEM) network (gem.tropicalforests.ox.ac.uk). Fieldwork was funded by grants from the UK Natural Environment Research Council (NE/I014705/1 and NE/P001092/1), European Research Council Advanced Investigator Grants to Y.M. (GEM-TRAITS: 321131), and an Africa Oxford Initiative Catalyst Grant to H.Z. ICP acknowledges support from the European Research Council (787203 REALM) under the European Union’s Horizon 2020 research programme. This work is a contribution to the LEMONTREE (Land Ecosystem Models based On New Theory, obseRvations and ExperimEnts) project, funded through the generosity of Eric and Wendy Schmidt by recommendation of the Schmidt Futures program. We also acknowledge the Wildlife Division of the Forestry Commission in Ghana, and Forest Research Institute of Ghana (FORIG) as well as the many field assistants who helped with data collection from the field.

## 6 Supplementary Method – Manual for reproducing results

The purpose of the supplementary material is to guarantee the reproducibility of all parameters used in our modelling work. As a result, it prioritizes technical details over readability, serving as a technical manual of the "Methods" chapter in the main text.

### 6.1 To reproduce Figure S3

(Codes in Zenodo deposit 1_prepare_climate_data and 3. supplementary_compare_climate_variable)

The ratio of leaf internal CO_2_ to eternal CO_2_ (c_i_/c_a_) was estimated from leaf δ^13^C measurements. We initially estimated the difference between the leaf stable isotope ratio and the atmospheric stable isotope ratio at that place and time (Δ^13^C) from δ^13^C, using the method described by a previous study ^77^. Subsequently, we calculated c_i_/c_a_ from Δ^13^C using equation 11 in a previous study ^46^. The community-weighted mean and standard error were then calculated based on 238 measurements. This isotope based c_i_/c_a_ was compared to Pmodel predicted c_i_/c_a_ explained in 6.2.2 Pmodel experiment (Figure S3).

The P-model requires several climate variables as input, and this can be a source of uncertainty in modelled GPP. To address this, we compared the temperature and relative humidity from local weather stations to products commonly used by vegetation models. We selected relative humidity and temperature provided by ERA-interim and shortwave radiation retrieved from ERA5-Land. However, the data-model differences we found for temperature, relative humidity and c_i_/c_a_ (calculated from temperature and vapour pressure) were very small and not likely to contribute significantly to the GPP data-model discrepancy (Figure S3). Moreover, data-model comparisons of many climate variables have been done in previous literature ^78^. Therefore, we used ERA-interim temperature, vapour pressure and calculated c_i_/c_a_ for all GPP experiments in this study to maintain consistency and ensure a fair comparison with other models (Figure 2), while the choice of PPFD data set was specific to the experiment (Figure 3).

For satellite based fAPAR, we extracted MODIS fAPAR (MOD15A2H) of the 14 plots from 2001 to 2020 and averaged them into mean fAPAR per site. To remove cloud contaminated fAPAR, we selected only CLOUDSTATE=00 and =10. Since MODIS GPP and fAPAR share the same CLOUDSTATE, this ensures that the fAPAR used in Pmodel_P is identical to that used in MODIS GPP (Figure 3). The issue of cloud contamination is visualized in Figure 1 and Figure 4. Field measured fAPAR is explained in 6.2.1.

### 6.2 To reproduce Figure 4 and Figure S2

(Codes in Zenodo deposit 4.Figure_two_GPP_from_measured_Vcmax)

This chapter explain how to calculate GPP for each Pmodel experiment as shown below.

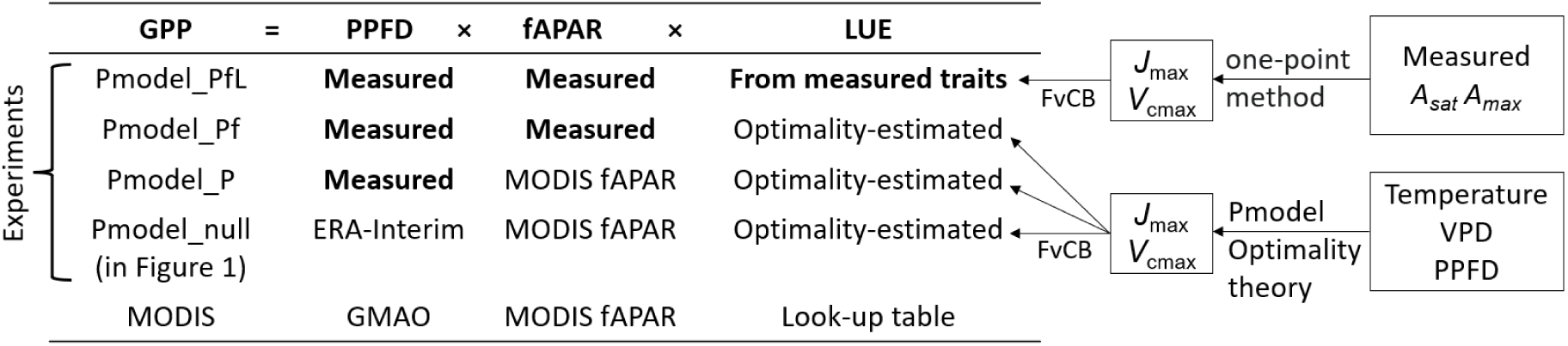

#### 6.2.1 Experiment Pmodel_PfL

We used measured mean fAPAR, light-saturated photosynthetic rate (*A*_sat_), light- and CO_2_- saturated photosynthetic rate (*A*_max_), χ, and climate variables for each site as inputs to the P-model in order to calculate ‘Pmodel_PfL GPP’ for each site. *V*_cmax_, *J*_max_ and LUE can also be calculated from the above measurements. We chose field-measured *A*_sat_ and *A*_max_ instead of field-measured *V*_cmax_ and *J*_max_ because the former is simpler to measure. We have multi-seasonal samplings and lots of replicates for *A*_sat_ and *A*_max_ (which do have strong seasonal variation) ^79^.

fAPAR was estimated using hemispherical photography. hemispherical images taken with a Nikon 5100 camera and Nikon Fisheye Converter FC-E8 0.21x JAPAN near the center of each of the 25 subplots in each plot in each site, at a standard height of 1 m, and during overcast conditions. 22,000 photos were collected in total, every month during 2016-2017(ANK), 2012-2017 (BOB&KOG). Photos were processed using machine learning-based software ‘ilastik’ ^29^ for pixel classification and CANEYE ^30^for leaf area index calculations. The exposure procedure followed ^80^ and GEM manual ^26^ (http://gem.tropicalforests.ox.ac.uk). The following parameters were supplied to CANEYE to calculate both leaf area index (LAI) and fAPAR

1. P1 = angle of view of the fish eye divided by the amount of pixels from centroid of the fish eye circle to where horizon is on the image.
2. angle of view = 90 degree, in which case, the edge of the photo is the horizon and the centroid of the image is zenith.
3. COI = 80, consideration of field is 80 degrees, we don’t want the edge of the photo because it is not clear and sometime obscure by tall grasses or saplings.
4. Sub sample factor =1
5. Fcover = 20 degree, this is to calculate the percentage of black pixels within central 20-degree ring. We used this to understand the relative openness of canopy for the given image. It is not relevant to LAI
6. PAIsat = 10, When a pixel is completely black, mathematically, the leaf area index (LAI) is infinite. As we provide CANEYE 25 subplot images for each estimation of LAI, this means all 25 subplot images show black at a given pixel. To address this ‘infinite’issue, we uses a value of 10 for LAI in such cases. This value is based on the guess that, the densest point in a tropical forest should had an LAI of 10.
7. Latitude 0 and Day of year a random number (not relevant for tropical site LAI) CANEYE reported a diffuse fAPAR and a direct fAPAR. We calculated overall fAPAR as 0.8*Diffuse_fapar +0.2*Direct_fapar ^81^. We used 0.8 and 0.2 because We assumed that 80% of days in the study region is cloudy.

To measure *A*_sat_ and *A*_max_, we used an open-flow gas exchange system (LI-6400XT, Li-Cor Inc., Lincoln, NE, USA). To ensure a proper representation of the forest stand, we sampled tree species that constituted approximately 80% of the plot basal area. For each species, we selected three mature and canopy emergent trees, and cut one fully sunlit and one shaded branch per tree using a single rope climbing technique. We immediately placed and recut the cut branch under water, and measured the maximum rate of net CO_2_ assimilation at 400 ppm CO_2_ (*A*_sat_) and 2000 ppm CO_2_ (*A*_max_) on three leaves per branch. The PPFD was set to 2000 μmol m^-2^ s^-1^ and block temperature was kept constant at 30 °C. Each measurement was derived into *V*_cmax_ and *J*_max_ at growth temperature ^46^ using R package ‘plantecophys’ ^31,32^. Measurements were made every three months from 2014 to 2016 to cover both wet and dry seasons. Although we measured both shade and sun leaves, we used sun leaves only in this study (consistent with common practice in field studies of photosynthetic traits). We used above -anopy PPFD as model input and we note that only sun leaves acclimate to this level of PPFD; shade leaves acclimate to darker environments and have consistently lower *A*_sat_ than sun leaves ^34^. We filtered outliers with the R package ‘outliers::scores’, ‘iqr’ method with threshold 1.5. Then measurements from 1394 leaves were used to calculate community-weighted means based on the basal area of each species. The same weights were applied in calculating standard errors ^82^.

**LUE was calculated from *A*_sat_ and *A*_max_ following the method below.**

We first **convert A_sat_ to V_cmax_** using ‘one-point’ method, equation 3 in ^32^.

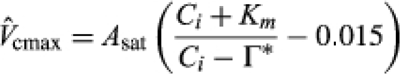

We used R package *Plantecophys* ^31^ to do so using the following codes.

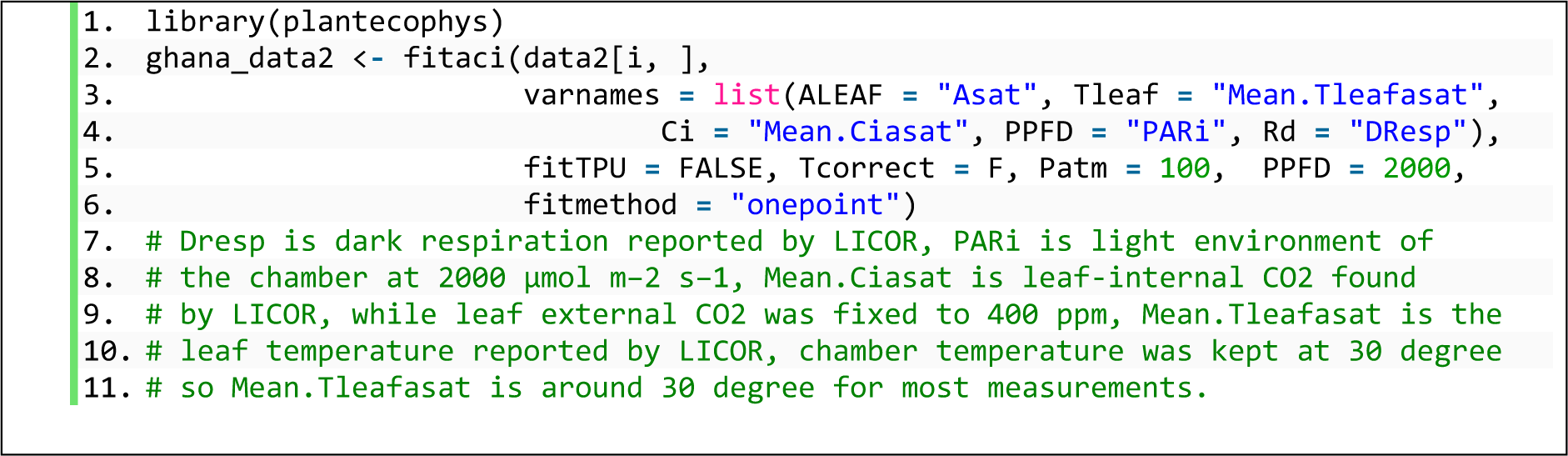

We then **convert *A*_max_ to J_max_**, using ‘Jmax limitation equation’.

However, there are two ‘J_max_ limitation methods’ that leads to two equations linking Amax and J_max_. Well, if you ignore ‘J_max_ limitation’ then simply *A*_max_ = J_max_ / 4. ^27^ explains the two equations very well. More information could be found from these papers ^43,44,83^.

In this paper, we used the first equation, which was equation 13 in study ^27^:

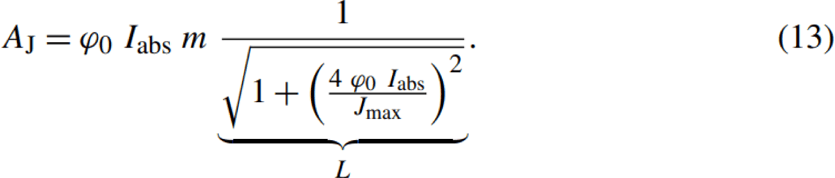

While there are also the second equation, which was equation F20 in study ^27^:

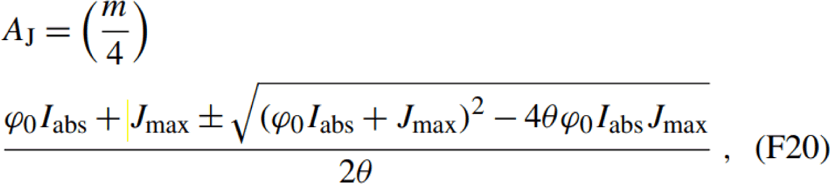

R package ‘*rpmodel’* (we will use this later) do have both equations written in, but for an unknown reason, results given by the second equation (called Smith19 in the package) have unreasonably low J_max_. It is more likely to be coding issue because the python version of ‘rpmodel’ reports reasonable results for both methods. The python version is called ‘pyrealm’ (https://pyrealm.readthedocs.io/en/latest/index.html) (https://pypi.org/project/pyrealm/). This is out of the scope of this paper, so results are not included. Eventually we used R package ‘rpmodel’ with the first equation (named ‘Wang17’ in the package).

However, package ‘*Plantecophys*‘ used the second equation. To be consistent with *rpmodel*, we modified source codes of *Plantecophys* in function *photosyn.R*, by **replacing**

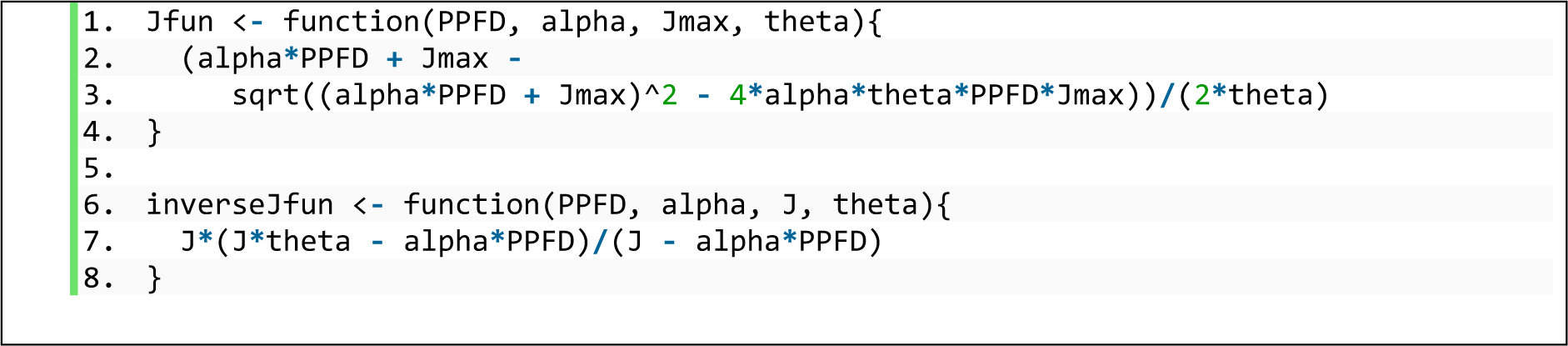

**With**

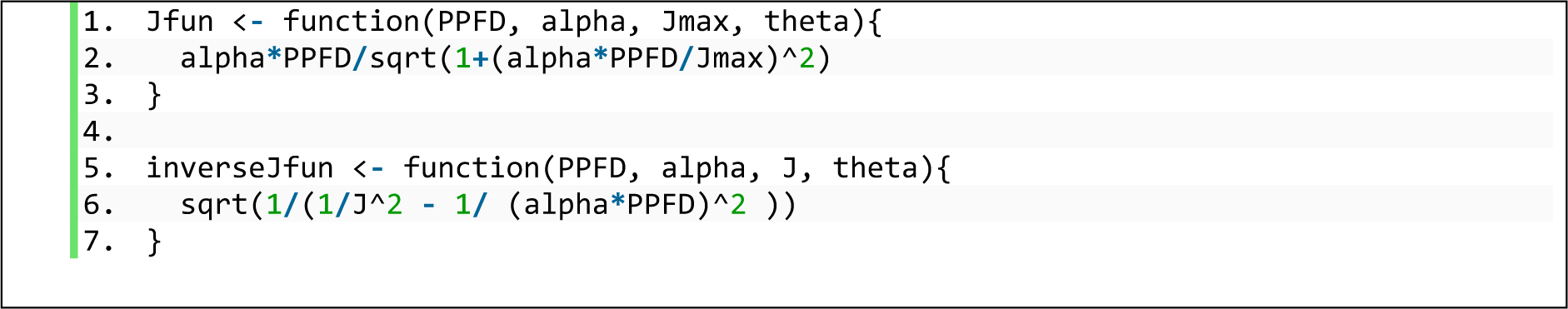

Now, J_max_ could be obtained by

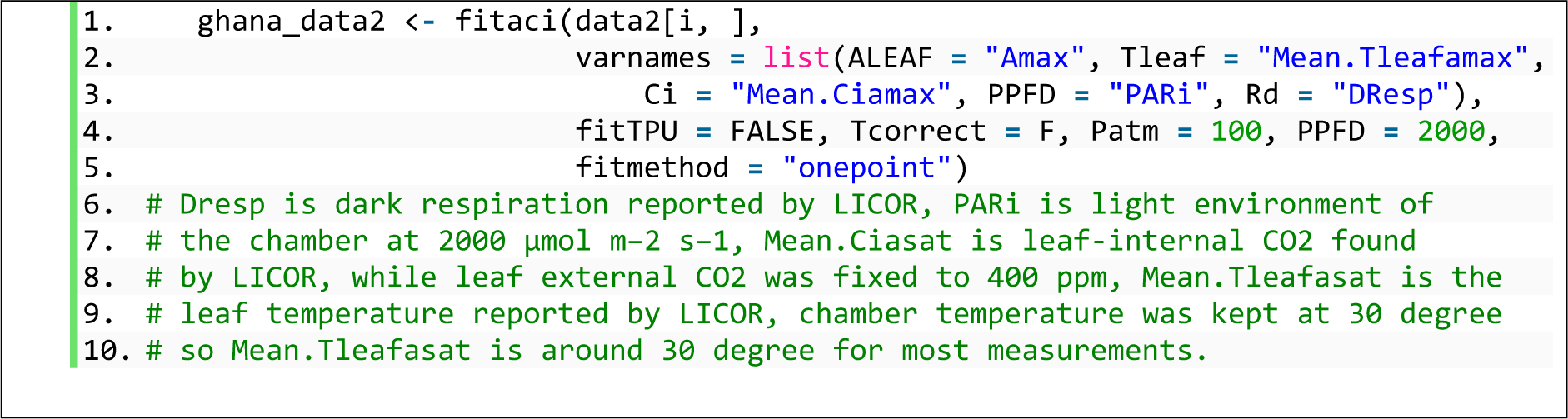

Note that temperature correction was turned off in the above. The outputs Vc_max_ and J_max_ are under measurement chamber temperature (around 30 degree). We used function ‘ftemp_inst_vcmax’ provided by *rpmodel* to standardise Vc_max_ and J_max_ from measurement temperature to mean annual air temperature of the site, using equation in ^27^. We tagged the above as Jmax_onepoint_tair and Vcmax_onepoint_tair.

Now, GPP, LUE, A_J_ and A_C_ could be calculated using Jmax_onepoint_tair and Vcmax_onepoint_tair.

First, A_C_ and A_J_ need to be calculated using:

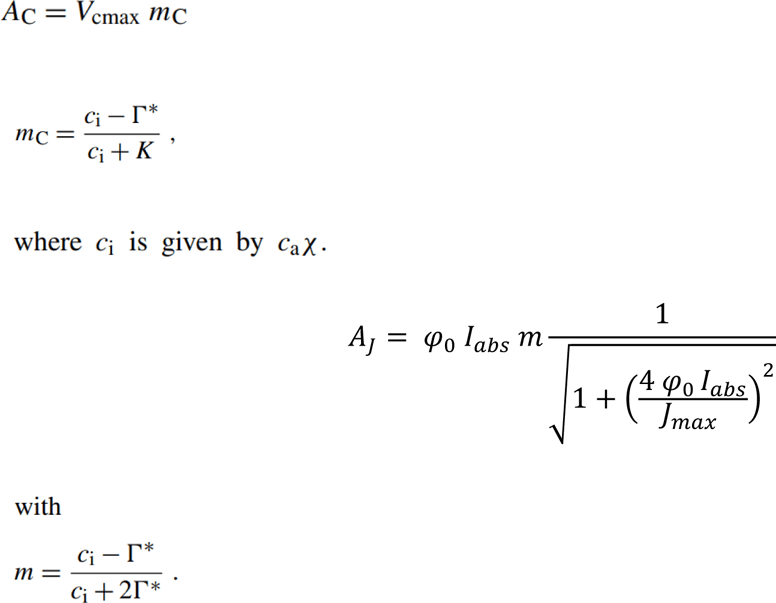

Next, we should check whether A_C_ = A_J_, following coordination hypothesis ^84^ (Figure S2). Upon checking coordination, we calculate the per unit leaf surface area assimilation rate following Aj only because (1) we are comparing to other light use efficiency model (MODIS) (2) Pmodel is essentially a light use efficiency model and was presented using Aj equation ^27,43^

The last step is to upscale per unit leaf surface area (*A*) to GPP using fAPAR. We could also calculate the traits-based LUE from A_C_, using: LUE = A_J_ / I_abs_.

Parameters involved above are available in ‘*rpmodel’* and explained in ^27^. The only difference is that we are now using the Jmax_onepoint_tair and Vcmax_onepoint_tair calculated above, instead of Pmodel prediction of optimality theory based Vc_max_ and J_max_.

The above could be done using codes in ‘*rpmodel’* specifically:

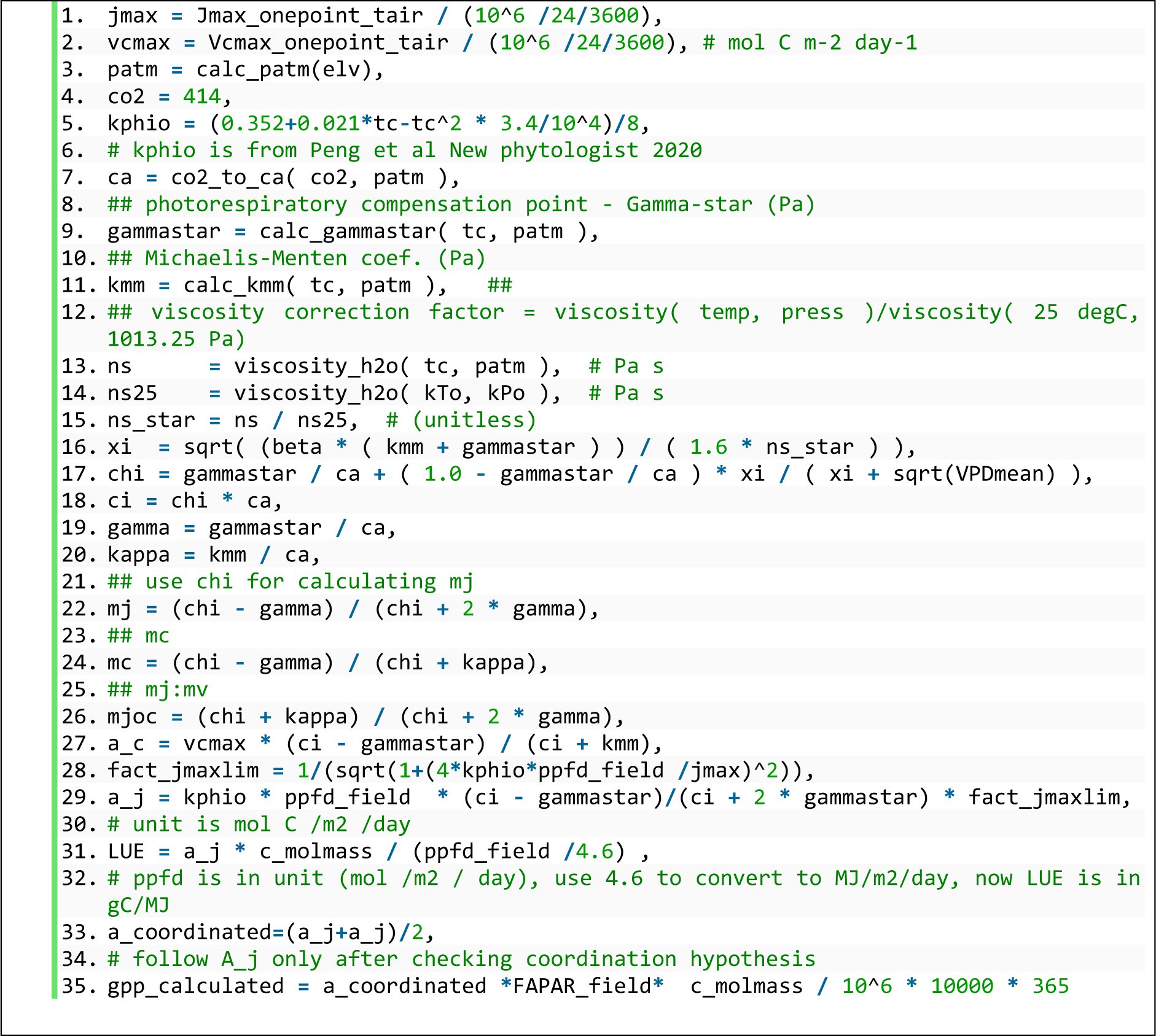

#### 6.2.2 Experiment Pmodel_null, Pmodel_P and Pmodel_Pf

The R package ‘rpmodel’ can be used to predict values of χ, *V*_cmax_ and *J*_max_ from climate variables. These values, combined with MODIS fAPAR (MOD15), were used to derive the Pmodel_P. Combined with field-measured fAPAR, they were used to derive Pmodel_Pf. Combined with MODIS fAPAR and ERA5 PPFD, they were used to provide Pmodel_0 (Figure 3).

Pmodel_null, Pmodel_P and Pmodel_Pf share the same codes as shown below, just with different input variables of fAPAR and PPFD (Figure 3).

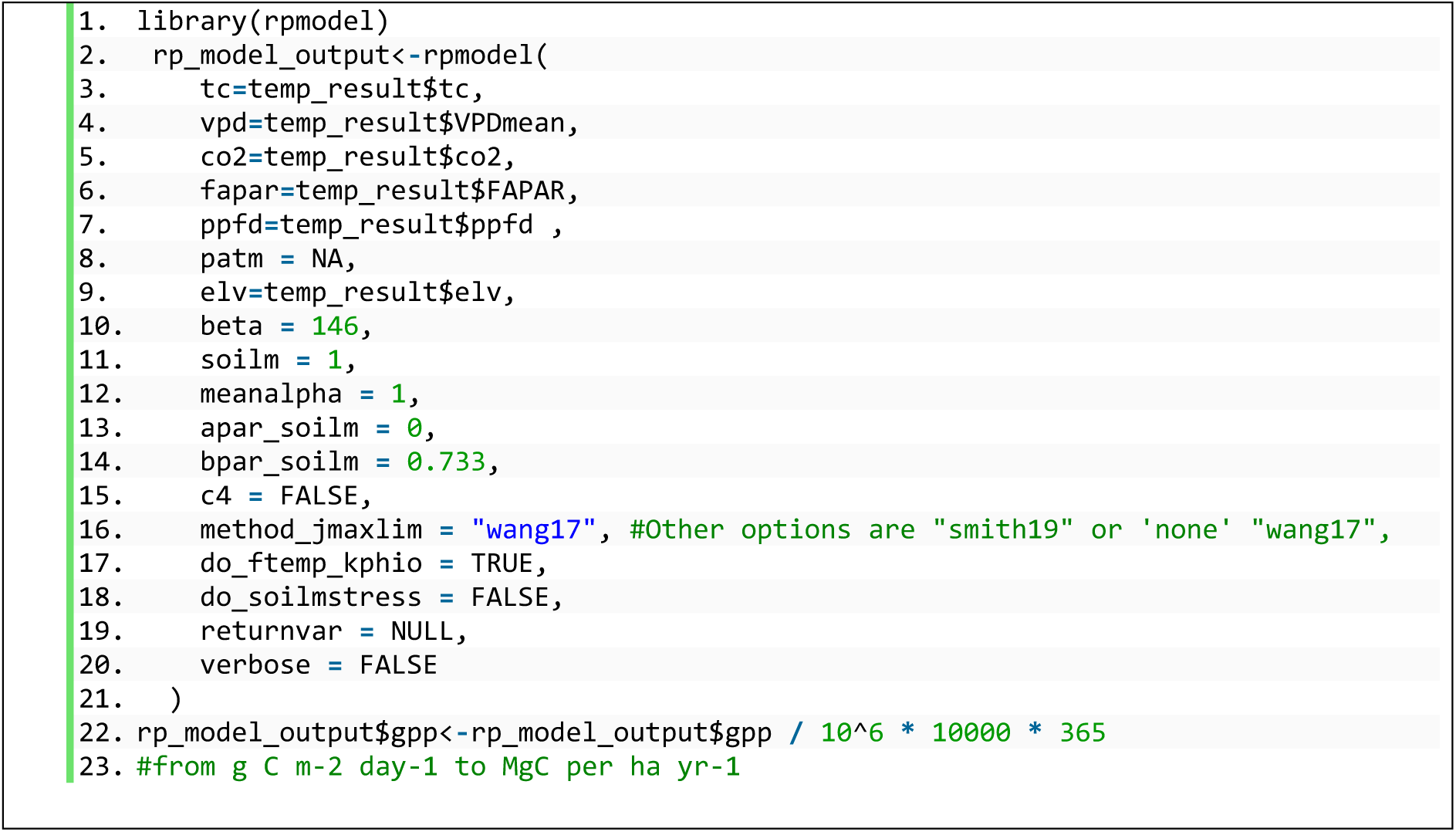

### 6.3 To reproduce Figure S1

Using Google Earth Engine, we retrieved MODIS GPP from the collection ‘MODIS/006/MOD17A3HGF’ for the period 20010101 to 20201231 (The gap filled and quality-controlled version of MOD17A2H); We drew a rectangular area of 0.5° x 0.5° (or 50 x 50 km) over Bobiri forest, where the study site BOB is located, and visualized the mean GPP of the area over the period (Figure S1).

### 6.4 To reproduce Figure 1 and Figure 4

(Codes in Zenodo deposit 2_Figure_four_timeseries_of_fapar)

For the period 20010101 to 20201231, we extracted MOD17A2H GPP of the 14 plots using their coordinates and calculated the mean GPP per site (not per plot). We also extracted MODIS fAPAR (MOD15A2H) of the 14 plots and averaged them per site. To remove cloud contaminated GPP and fAPAR, we selected only CLOUDSTATE=00 and =10. Since MODIS GPP and fAPAR share the same CLOUDSTATE, this ensures that the fAPAR used in Pmodel_PPFD is identical to that used in MODIS GPP (Figure 3) (see Rationale).

To illustrate the spatial cover of fAPAR cloud contamination we used MODIS MOD15A2H at timestamp A2016249. Pixels contaminated by cloud were identified using ‘cloudstate = 01’ provided in band ‘faparlaiQC’ (Figure 1). To illustrate the seasonal variation of fAPAR and cloud status, we calculate monthly average from 20010101 to 20201231 for fapar and for the percentage of cloud contamination (cloudstate = 01) (Figure 5).

### 6.5 To reproduce Figure 2 and Figure S4

(Codes in Zenodo deposit 5.compare_to_trendy_flux_com)

MODIS GPP was obtained from MOD15A2H as explained above (See 6.4).

There is a flux tower at ANK (GH-Ank), but not at BOB or KOG. Therefore, we compared biometric GPP at the three sites with FLUXCOM. For FLUXCOM, we chose the RS_METEO version instead of RS because the magnitude of GPP in RS_METEO does not involve uncertainty from MODIS FAPAR, which makes the comparison between FLUXCOM and MODIS GPP more independent. Note that the seasonal variation of GPP in RS_METEO version does involve uncertainty due to MODIS fAPAR, but this seasonal variation was not studied here. For TRENDY, we analysed the model outputs in version 9 under the S2, in which climate and CO_2_ change while land use is kept constant. We extracted values of the grid cells where the three study sites stand, for output "gpp," "gpppft," and "landcoverfrac." The latter two together could determine a forest-only GPP (Figure S4), where we grouped land use types including ‘evergreen forests’, ‘deciduous forests’ or ‘raingreen forests’ etc. For both TRENDY and FLUXCOM, we extracted the grid cells where the forest plots fall, as an average from 2009 to 2019.

## 7 Supplementary figures

**Figure S 1.**
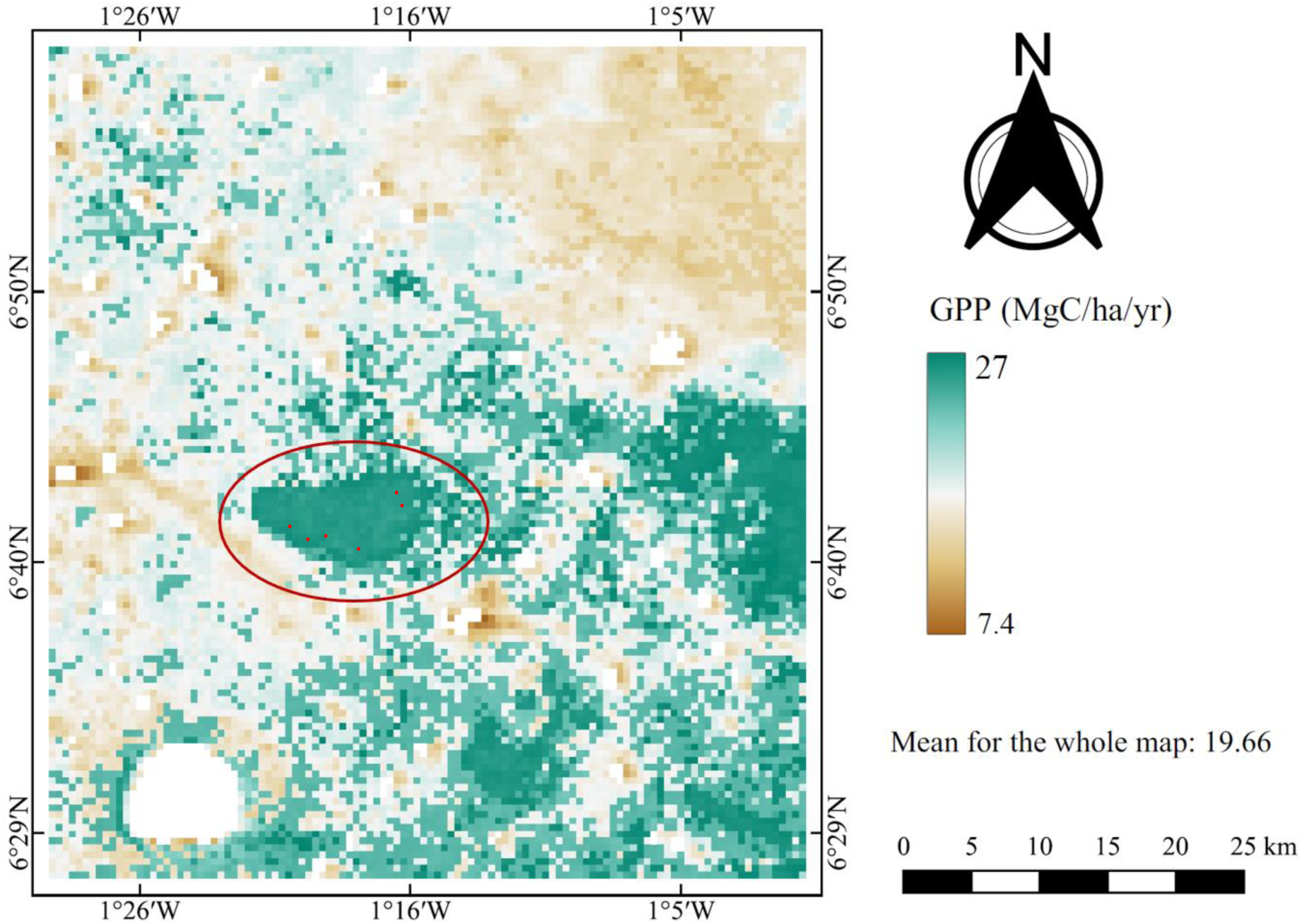
MODIS gross primary productivity (GPP). The red circle highlights Bobiri (BOB) forest reserve, surrounded by farmland. Six red dots denote six plots (100 x 100 m) established for GPP measurements in this forest reserve. The whole map area is roughly equal to a grid cell in TRENDY models, the typical resolution of which is half-degree (or 50 x 50 km). We visualized the GPP of the area averaged over the period 20010101 to 20201231 using product MOD17A3HGF (500m resolution, much finer than TRENDY), which has been cloud contamination-filtered and gap-filled. The figure shows that (1) West African forests are very fragmented and the study site BOB is surrounded by farmland, which makes plant functional type recognition an important issue in modelling (see hypotheses for Objective 3 in main text). (2) MODIS GPP and TRENDY models are on different resolution. Study plots fall into different pixels of MODIS but into the same pixel of a carbon model.

**Figure S 2.**
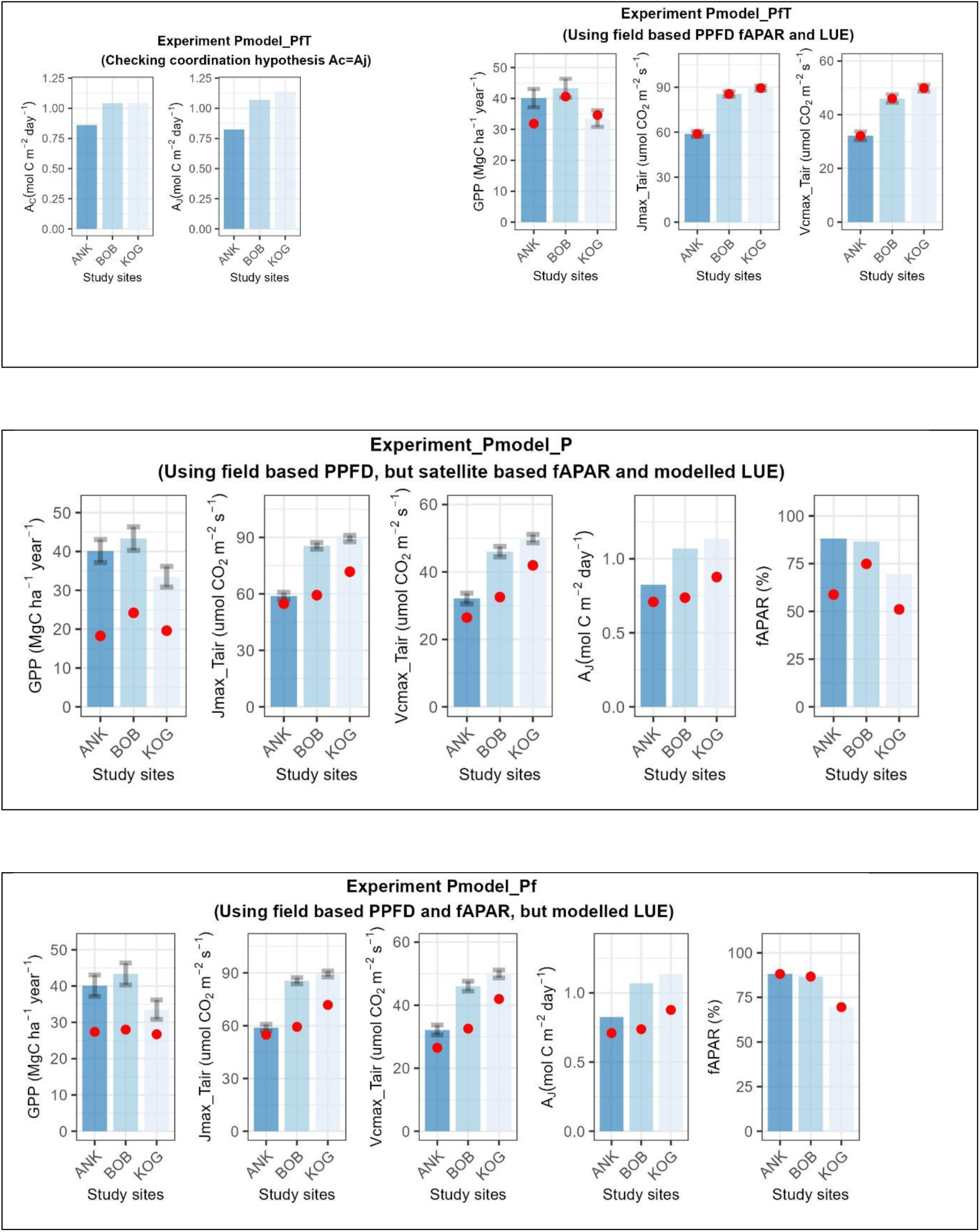
Comparing biometric gross primary productivity (GPP, MgC/ha/year) (Blue bar) to GPP of multiple experiments (red dots). Precipitation decreases from ANK to BOB, KOG. The figures also show data (blue bars) model (red dots) comparison for Rubisco carboxylation capacity (Vcmax_Tair, umol CO_2_ m^-2^ s^-1^), and electron transport capacity (Jmax_Tair, umol CO_2_ m^-2^ s^-1^) under mean annual air temperature, photosynthesis on the enzyme-limited (A_C_, mol C m^-2^ day^-1^) and electron transport-limited (A_J_, mol C m^-2^ day^-1^) rates under mean annual air temperature. Note that for experiments Pmodel_P and Pmodel_Pf, Pmodel predicts Vcmax and Jmax with the assumption that A_C_ = A_J_ (coordination hypothesis). Therefore, only A_J_ was displayed. For experiment Pmodel_PfT however, A_C_ and A_J_ are independently calculated and thus A_C_ is not identical to A_J_. All variables were calculated under mean annual air temperature not standardized to 25 °C.

**Figure S 3.**
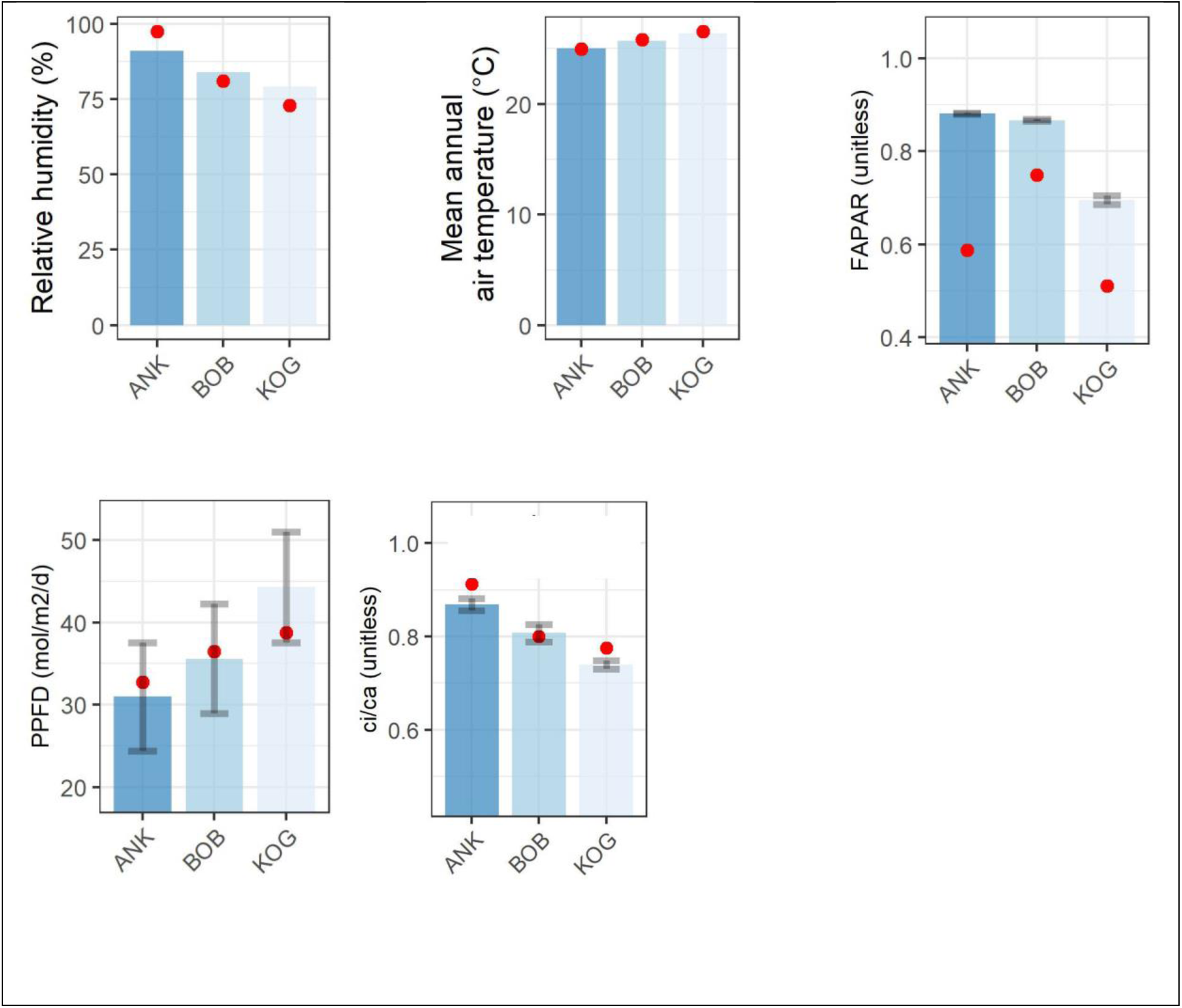
Field measurements (blue bars) and model (or from global products, red dots) comparison for relative humidity, mean annual temperature, leaf internal to external CO_2_ concentration (c_i_/c_a_), fraction of absorbed photosynthetically active radiation (fAPAR) and photosynthetic photon flux density (PPFD). See Supplementary notes for source of data.

**Figure S 4.**
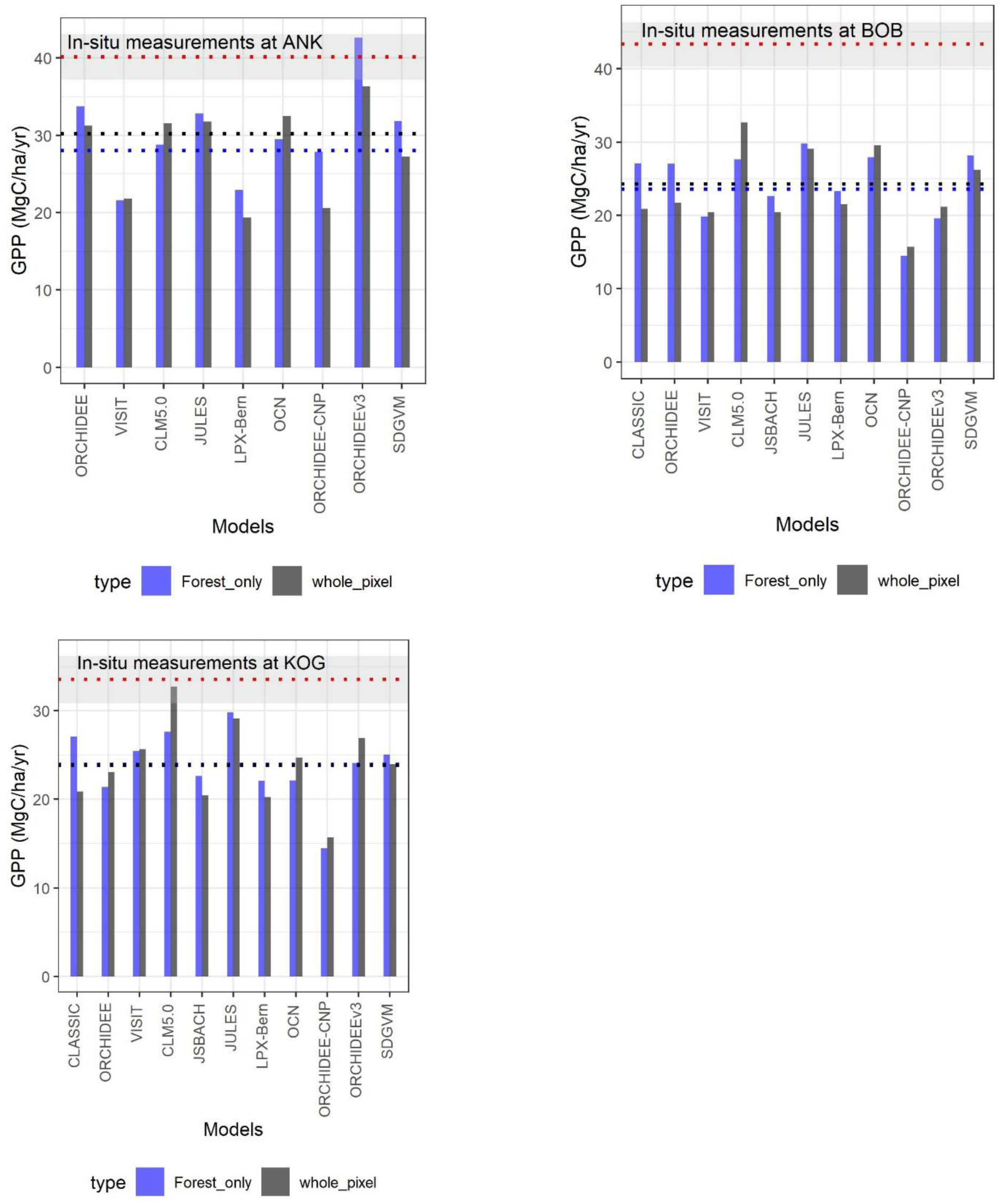
Comparing Forest-only gross primary productivity (GPP) (blue) to the GPP of the whole pixel (black), which is a mixture of agricultural land, forest and grass at the three study sites, with percentage vary considerably among models. The analysis is only possible for TRENDY models that reported GPP of each plant functional type. The red line and grey zone denote field measured GPP and its uncertainty. Figure 2 has more models than Figure S 4 because some models do not report GPP per plant functional type. Site ANK has less models than BOB and KOG because ANK is near the coast and the pixel was classified as ‘ocean’ in some models. The readme file of LPJ-GUESS asks not to upscale GPP (variable gpp_pft) based on land cover fraction (variable landCoverFrac), because of which LPG-GUESS was neglected.

**Table S 1.**
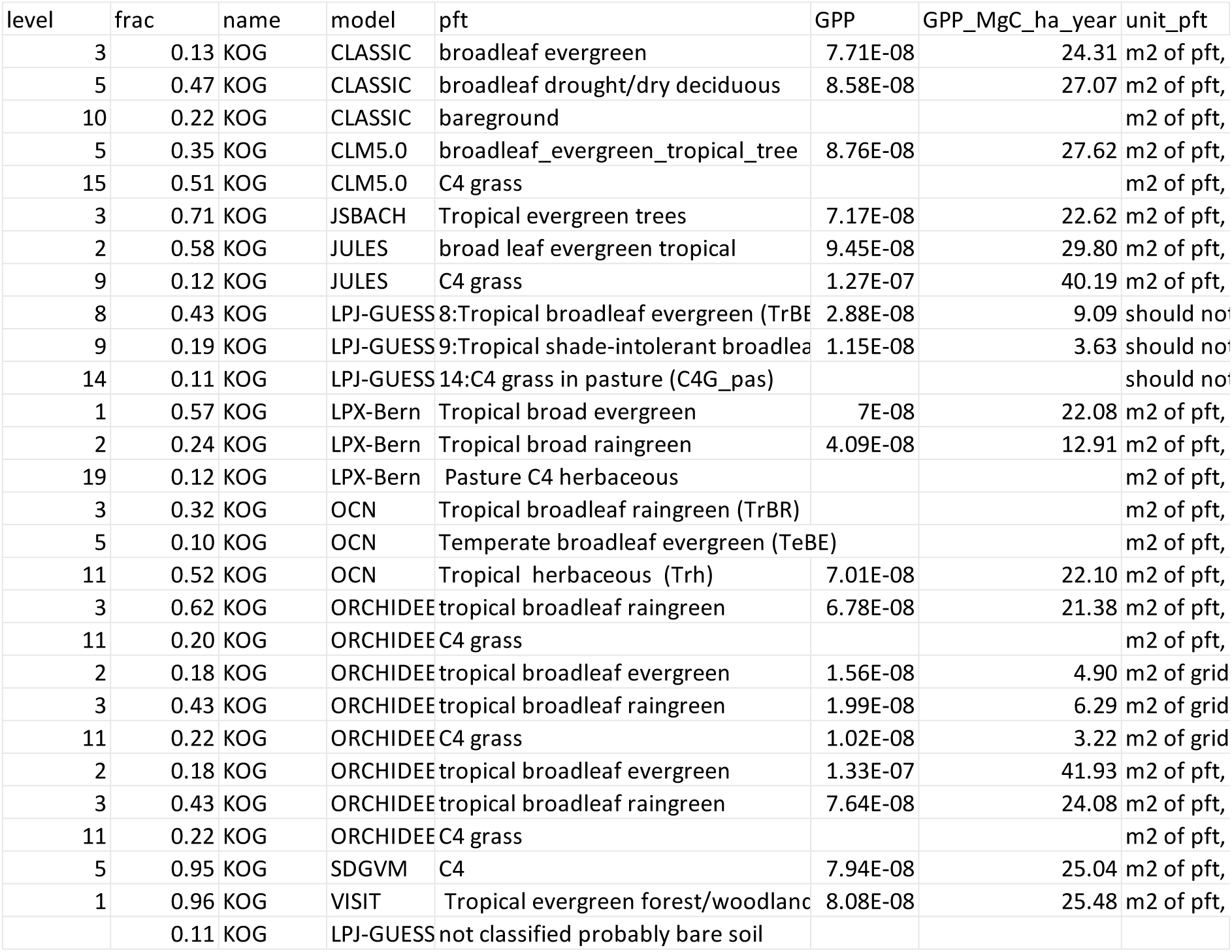
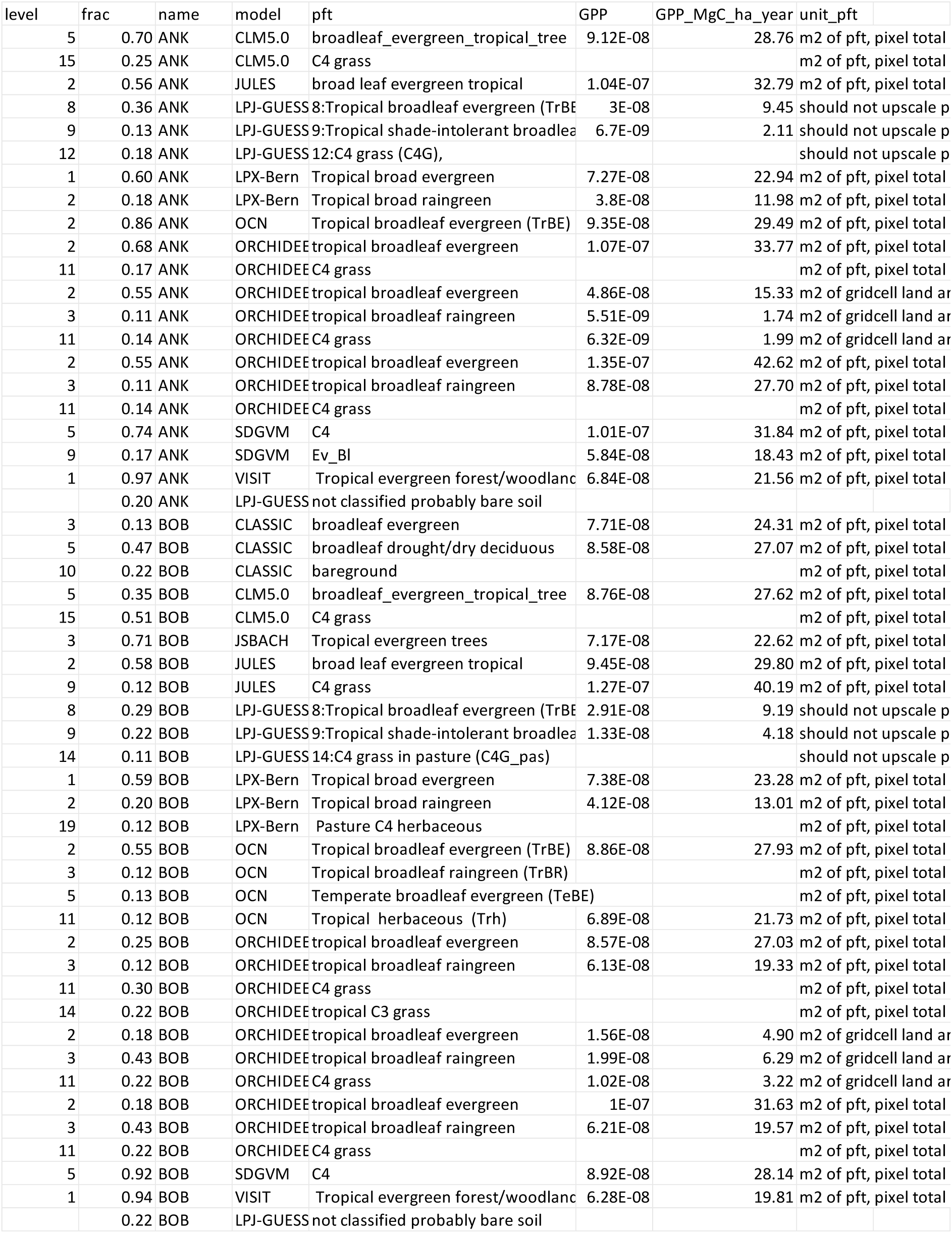
Extracting variable ‘gpppft’ and variable ‘landcoverfrac’ from TRENDY models for each pixel where our study sites fall into. Longitude and latitude are coordinates of the study sites. ‘Name’ is the name of study site. GPP is the value contained in variable ‘gpppft’ in original unit of model outputs. Models normally have multiple plant functional type. Each has a unique ID. ‘Level’ is the id. GPP_MgC_ha_year is converted from column GPP, standardised to unit MgC_ha_year. ‘Frac’ is the percentage fraction of such plant functional type in such model at the grid cell. ‘pft’ the name of plant functional type defined by the model. ‘Unit’ explains how to link GPP_MgC_ha_year here (stored in variable gpp_pft) with total GPP of a grid cell (variable gpp). For most models, you need to multiply "frac" first, but for some model (e.g. LPJ-GUESS), the "multiplied with frac" has been done by the modeler so you just need to sum all gpp-pft records under this model. Note: frac < 10% was neglected. Model ISAM was neglected because we could not find plant functional type information in their nc files. PFT noted as ‘not classified probably bare soil’ implies that frac of the pixel does not sum up to 1, the remaining was assumed bare soil as told by LPJ-GUESS user manual. **Only a screenshot of part of the table is shown below, please download the full table from supplementary data.**

**Table S 2.** Comparing rubisco carboxylation capacity (Vcmax, umol CO_2_ m2 s-1) at 25 °C for various plant functional types (PFT) among (1) DGVMs, provided by ^62^ (2) derived from field measured *A*_sat_ and *A*_max_, same as Figure S2 but standardised to 25 °C, and (3) Pmodel_P experiment, same as Figure S2 but standardised to 25 °C. Note that modelled values are compared to field measurements derived values where arbitrary colours are given; green denotes a similar value. Red denotes that the model severely over-estimates while blue denotes that the model severely underestimates V_cmax_ for West African forests. Note that the actual Vcmax used in the model for West African forests could be different to the values presented here because the values are recalculated for the entire PFT ^62^; The actual Vcmax could vary within a PFT depending on the model; The values recalculated by ^62^ could also be different to others depending on the specific models version and simulation scenarios, for examples, comparing BETHY to ^63^. It is not possible to retrieve Vcmax directly from TRENDY simulation and thus, Table S1 and Table S2 are not mechanistically associated. Models listed here could be a sub-model or predecessor of TRENDY models (listed in Table S1). For example, BETHY is a sub-model of JSBACH ^63^; CTEM is a predecessor of CLASSIC ^85^. C4 grass is also presented because many models tag a portion of PFT of the study sites as C4 grass.

| Model | Plant functional type | $V_{cmax25}$ ($\mu\text{mol m}^{-2} \text{s}^{-1}$) |
| --- | --- | --- |
| Field measured as shown in (Figure S2) but standardised to 25 °C | ANK(broadleaf evergreen) | 30.49 |
| Field measured as shown in (Figure S2) but standardised to 25 °C | BOB(mixture of broadleaf evergreen and raingreen) | 39.07 |
| Field measured as shown in (Figure S2) but standardised to 25 °C | KOG(broadleaf raingreen) | 39.61 |
| Pmodel_P as shown in (Figure S2) but standardised to 25 °C | ANK(broadleaf evergreen) | 26.54 |
| Pmodel_P as shown in (Figure S2) but standardised to 25 °C | BOB(broadleaf evergreen+raingreen) | 30.35 |
| Pmodel_P as shown in (Figure S2) but standardised to 25 °C | KOG(broadleaf raingreen) | 36.55 |
| AVIM | Broad leaf evergreen tropical tree | 64 |
| AVIM | Broad leaf deciduous tropical tree | 56 |
| AVIM | C 4 grass | 25 |
| BETHY | Tropical tree (oxisols) | 28 |
| BETHY | Tropical tree (non-oxisols) | 36 |
| BETHY | C 4 grass | 20 |
| Biome-BGC | Evergreen broad leaf forest | 51 |
| Biome-BGC | Deciduous broad leaf forest | 47 |
| Biome-BGC | C 4 grass | 60 |
| CLM | Broad leaf evergreen tree—tropical | 59 |
| CLM | Broad leaf deciduous tree—tropical | 34 |
| CLM | C 4 grass | 33 |
| CTEM | Broad leaf evergreen | 35 |
| CTEM | Broad leaf deciduous—dry | 65 |
| CTEM | C 4 grass | 30 |
| Hybrid | Grassland | 20 |
| Hybrid | Deciduous | 31 |
| Hybrid | Evergreen | 43 |
| Hybrid | Rain forest | 49 |
| IBIS | Tropical broad leaf evergreen | 163 |
| IBIS | Tropical broad leaf drought-deciduous | 163 |
| IBIS | Warm grass | 38 |
| JULES | Broad leaf tree | 37 |
| JULES | C 4 grass | 24 |
| O–CN | Tropical broad leaved evergreen | 24 |
| O–CN | Tropical broad leaved rain green | 20 |
| O–CN | C 4 herbaceous | 13 |
| ORCHIDEE | Tropical broad leaf evergreen trees | 18 |
| ORCHIDEE | Tropical broad leaf rain green trees | 22 |
| ORCHIDEE | Natural C 4 grass | 28 |

## 8 Supplementary information – Description of the study sites

The following study sites description information is also available at ^21^.

The study focuses on an aridity gradient established in Ghana, West Africa (Figure 4), that features a strong variation in aridity with only a 1.4 °C change in mean annual temperature (Table S3). The gradient contains three wet (mean annual precipitation, MAP 2000 mm) evergreen forest one-hectare plots in the Ankasa national forest (ANK-01 to ANK-03) in the extreme southwest of Ghana, close to the Côte d’Ivoire border. Two hundred kilometres to the northeast, there are six semideciduous one-hectare forest plots (MAP 1500 mm) located within Bobiri Forest Reserve (BOB-01 to BOB-06), close to the second largest city in Ghana, Kumasi. An additional one hundred kilometres to the northeast, there are six one-hectare plots located within the Kogyae Strict Nature Reserve, which encompasses the dry forest (MAP 1200 mm) to savanna transition zone (KOG-01 to KOG-06). All these plots are part of the African Tropical Forest Observation Network (AfriTRON) and ForestPlots.

The sites were at least 100 km away from each other. Plots within the same site were approximately 1 km away from each other. Therefore, the three study sites were in different grid cells of a model (or satellite product), but plots belonging to the same site were in the same grid cell. This study is thus conducted on site scale.

### KOGYAE (dry, including KOG-01 to KOG-06)

The Kogyae Strict Wildlife Reserve (7.28464, -1.42414) is located near the Ejisu-Juaben District of the Ashanti Region, Ghana. The reserve covers 386 km2 and was created in 1971. From KOG01 to KOG06, it spans from semideciduous forest to open grassland (forest-savanna transition). In the past, the forest was a continuous block that covered the whole reserve’s centre and took up nearly one-third of its space. However, during the 1980s and the early 1990s, logging, farming, and bushfires all contributed to a significant loss of forest cover and the consequent fragmentation of the forest into isolated areas. The main wet season lasts from April through June with a secondary wet season from September to November, which in total delivers 1200-1300 mm average annual rainfall. From April to June, Kogyae experiences its primary wet season, with a secondary season occurring between September and November. The total average annual rainfall received during these seasons is approximately 1200-1300 mm. The Voltarian system characterises Kogyae’s geology, and exposed rocks are typically reddish-brown sandstone. The soil in the savanna and transition regions is Haplic Arenosols, with thin, sandy loam topsoil, while the forest sites have Haplic Nitosols, which are more acidic than the savanna soil ^25,49^.

### BOBIRI (middle, including BOB-01 to BOB-06)

The Bobiri Forest Reserve (Lat 6.68697, Long -1.34401) covers 54.65 km2 and was established in 1931. It is largely covered with semideciduous and old-growth forest, which has a multilayered canopy structure, including an understorey layer composed of both shade-tolerant and shade-intolerant species, large coarse woody debris in all decay stages on the forest floor, and the presence of ferns. This site has six 1 ha plots (BOB-01 to BOB-06) with different levels of logging: BOB-01 and BOB-02 were last logged approximately 50 years ago (baseline conditions plots); BOB-05 and BOB-06 were last logged in 2012. The 10-yr annual rainfall ranges from 1210 to 1800 mm, with a dry season lasting from December to mid-March. The soils are Oxisols, deeply weathered, and highly acidic (3.5–4.0 in pH).

### ANKASA (wet, including ANK-01 to ANK-03)

The Ankasa National Forest (Lat 5.26313, Long -2.57924) was created in 1976 and has an area of 500 km2. The reserve is mostly covered by tropical evergreen rainforest. The mean annual precipitation (MAP) is approximately 2,000 mm, mainly concentrated from March to mid-July and from September to November. A dry period extends from December to February. The relative humidity is high throughout the year; daily, it ranges from 90% at night to 75% in the early afternoon. The landscape is characterised by the presence of small, gentle hills with an average elevation of 90 m.a.s.l., such that rugged and deeply divided terrains characterise the forest, with ANK03 in the swampy valley and others on the drier hills. The deeply weathered soils are highly acidic, 3.5–4.0 in pH, and on a broad basis classified as Oxisols.

**Table S3.** Study plots information. These are all one-hectare plots. Data source from ^21^

| Plot_code | RH (%) | vwc (%) | MCWD (mm) | MAT (°C) | MAP (mm) | Lat | Lon |
| --- | --- | --- | --- | --- | --- | --- | --- |
| ANK-03 | 91 | 11.63 | -13 | 25 | 2050 | 5.27102 | -2.69234 |
| ANK-02 | 91 | 6.78 | -13 | 25 | 2050 | 5.268485 | -2.695035 |
| ANK-01 | 91 | 5.96 | -13 | 25 | 2050 | 5.267868 | -2.693635 |
| BOB-03 | 83.9 | 11.37 | -374 | 25.7 | 1500 | 6.694531 | -1.293695 |
| BOB-04 | 83.9 | 9.42 | -374 | 25.7 | 1500 | 6.69096 | -1.317038 |
| BOB-05 | 83.9 | 8.64 | -374 | 25.7 | 1500 | 6.692606 | -1.30727 |
| BOB-06 | 83.9 | 8.19 | -374 | 25.7 | 1500 | 6.691368 | -1.307001 |
| BOB-02 | 83.9 | 7.87 | -374 | 25.7 | 1500 | 6.69147 | -1.338429 |
| BOB-01 | 83.9 | 6.4 | -374 | 25.7 | 1500 | 6.704713 | -1.319068 |
| KOG-02 | 79.2 | 4.13 | -412 | 26.4 | 1200 | 7.262316 | -1.149953 |
| KOG-03 | 79.2 | 2.9 | -412 | 26.4 | 1200 | 7.306792 | -1.156446 |
| KOG-05 | 79.2 | 2.54 | -412 | 26.4 | 1200 | 7.305341 | -1.164546 |
| KOG-04 | 79.2 | 2.42 | -412 | 26.4 | 1200 | 7.302644 | -1.180213 |
| KOG-06 | 79.2 | 1.72 | -412 | 26.4 | 1200 | 7.329423 | -1.15578 |

Note: (1) Volumetric water content (vwc) at site ANK, BOB and KOG are measurements of topsoil (12 cm) only. This data is provided for reference only and are not suggested for any quantitative analysis. (2) RH = relative humidity; vwc= volumetric water content, indicating surface soil moisture; MCWD = maximum climatological water deficit; MAT= mean annual air temperature; MAP = mean annual air precipitation; Lat = latitude; Lon = longitude. Plots are ranked by MCWD.

Table S3 (continue)
| Plot | Elev | PPFD | Trees | P | N | C | Ca | K | Mg | Sand | Clay | Silt |
| --- | --- | --- | --- | --- | --- | --- | --- | --- | --- | --- | --- | --- |
|  | (m) | (mol/m<sup>2</sup>/day) | (#/ha) | (mg/kg) | (%) | (%) | (mg/kg) | (mg/kg) | (mg/kg) | (%) | (%) | (%) |
| ANK-03 | 86 | 30.97 | 517 | 109.7 | 0.12 | 1.9 | 40 | 33.7 | 29.2 | 75.9 | 12.8 | 11.4 |
| ANK-02 | 124 | 30.97 | 445 |  |  |  |  |  |  |  |  | 15.3 |
| ANK-01 | 114 | 30.97 | 476 | 146.8 | 0.17 | 2.6 | 26.8 | 32.3 | 42 | 63.1 | 21.6 | 15.3 |
| BOB-03 | 294 | 35.58 | 631 |  |  |  |  |  |  |  |  |  |
| BOB-04 | 272 | 35.58 | 506 |  |  |  |  |  |  |  |  |  |
| BOB-05 | 246 | 35.58 | 527 |  |  |  |  |  |  |  |  |  |
| BOB-06 | 278 | 35.58 | 545 |  |  |  |  |  |  |  |  |  |
| BOB-02 | 281 | 35.58 | 789 | 258.3 | 0.16 | 1.7 | 657.6 | 49 | 133.7 | 46.7 | 28.8 | 24.5 |
| BOB-01 | 277 | 35.58 | 519 | 77.8 | 0.09 | 0.8 | 306.3 | 47.6 | 79.7 | 64.2 | 6.7 | 26.8 |
| KOG-02 | 229 | 44.26 | 197 | 67.2 | 0.06 | 0.7 | 378.9 | 42.5 | 75.6 | 82.4 | 2.3 | 15.34 |
| KOG-03 | 198 | 44.26 | 216 |  |  |  |  |  |  |  |  |  |
| KOG-05 | 221 | 44.26 | 193 | 81.9 | 0.04 | 0.6 | 237.1 | 28.7 | 81.3 | 76.9 | 4.3 | 18.71 |
| KOG-04 | 230 | 44.26 | 234 | 74.6 | 0.05 | 0.7 | 308 | 35.6 | 78.7 | 79.7 | 3.3 | 17.02 |
| KOG-06 | 195 | 44.26 | 202 |  |  |  |  |  |  |  |  |  |

Note: Plots are ranked by MCWD; Empty cells are due to the lack of data. Elev, elevation; MAP, mean annual precipitation; PPFD, photosynthetic photon flux density, calculated from shortwave radiation \*0.45; Trees, total number of trees larger than 10cm diameter at breast height, as this number change from year to year, the first year is picked if a plot has multiple years censuses; Soil nutrients (P, phosphorus; N, nitrogen; C, carbon; Ca, calcium; Mg, magnesium), and soil percentage of sand (Sand) and of clay (Clay)

## Ankasa

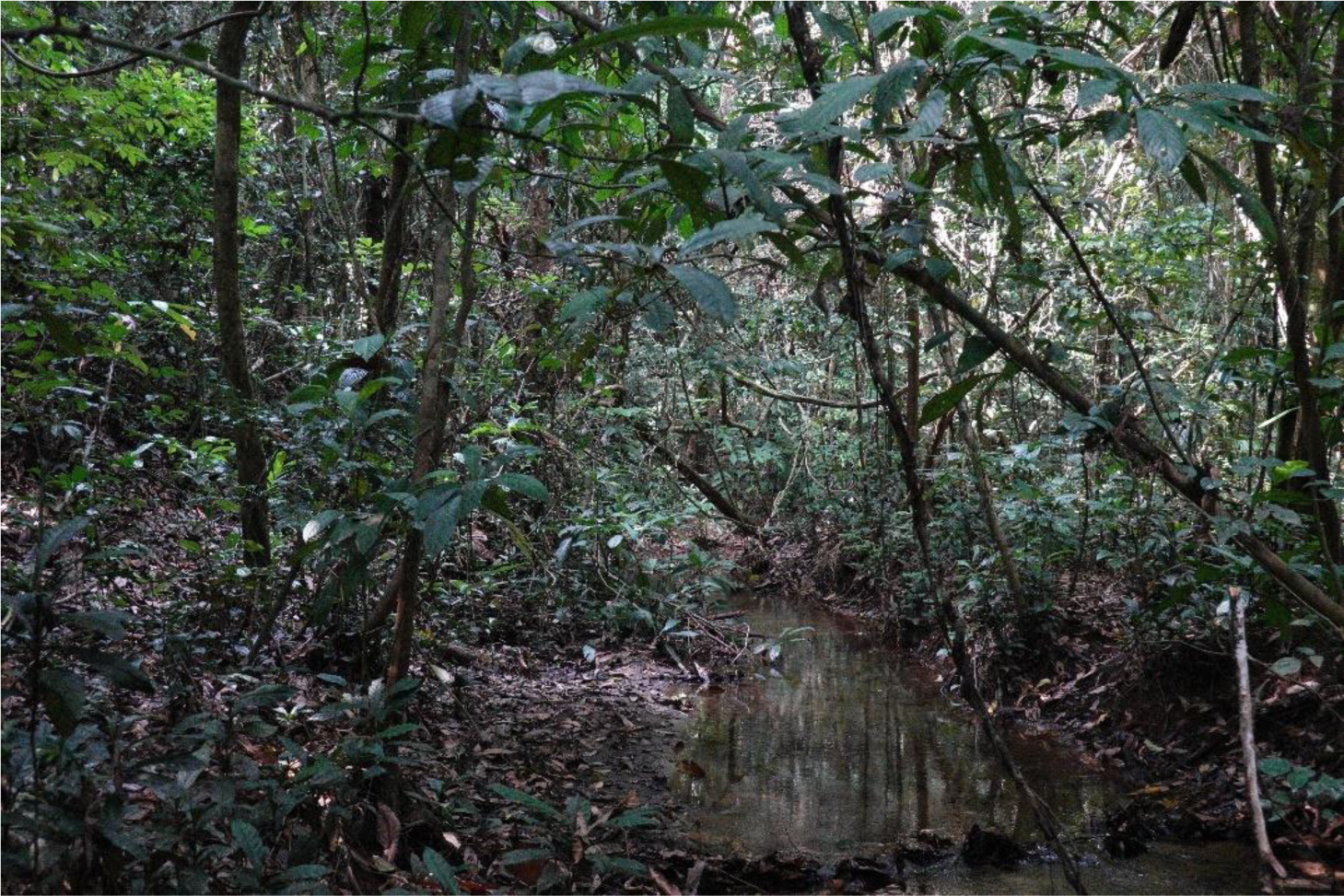

There is a stream running through ANK03 which largely floods the plot in the wet season. Ankasa - ANK01

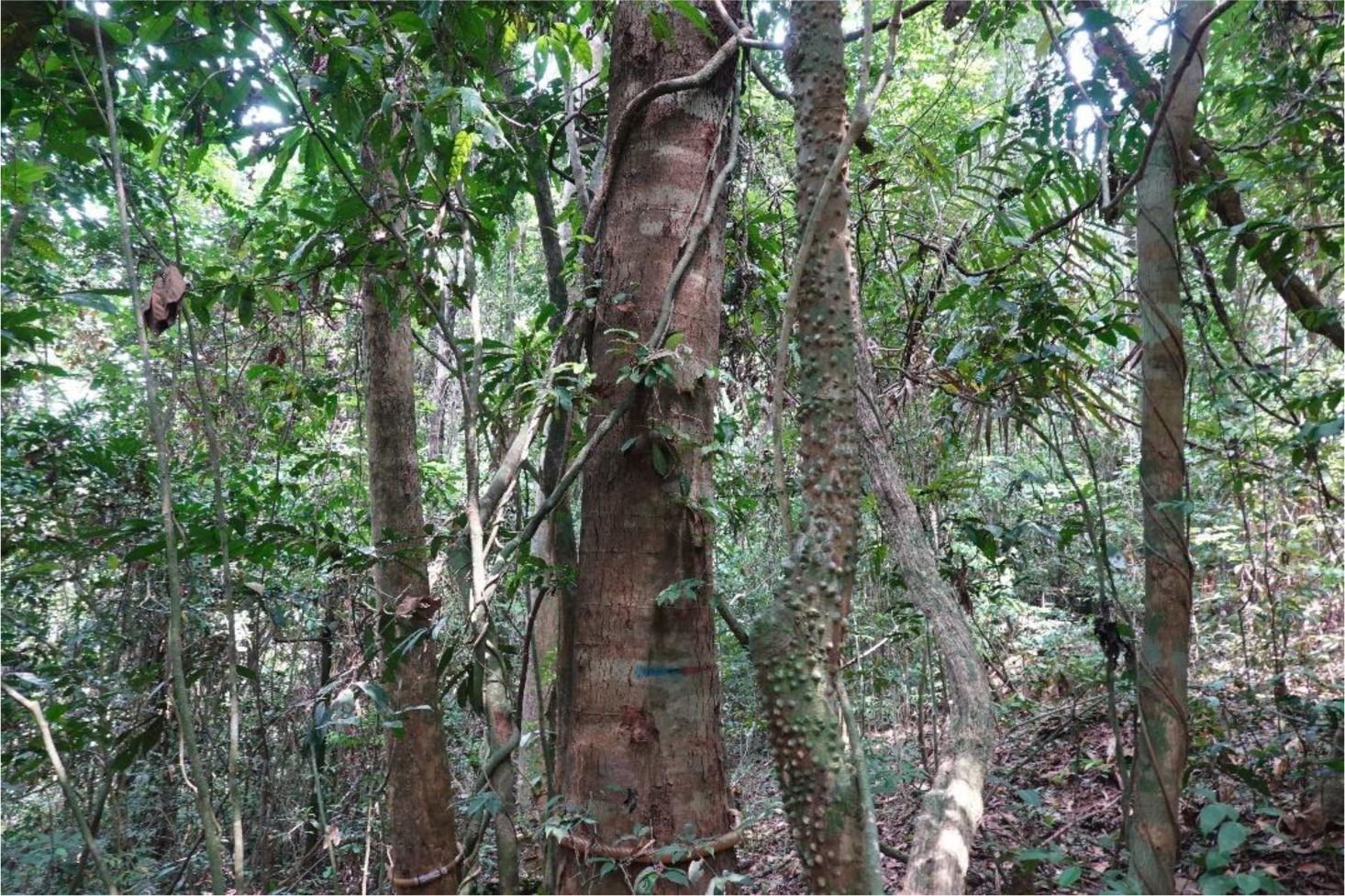

ANK01 and ANK02 are located on well-drained local hilltops.

## Bobiri - BOB01

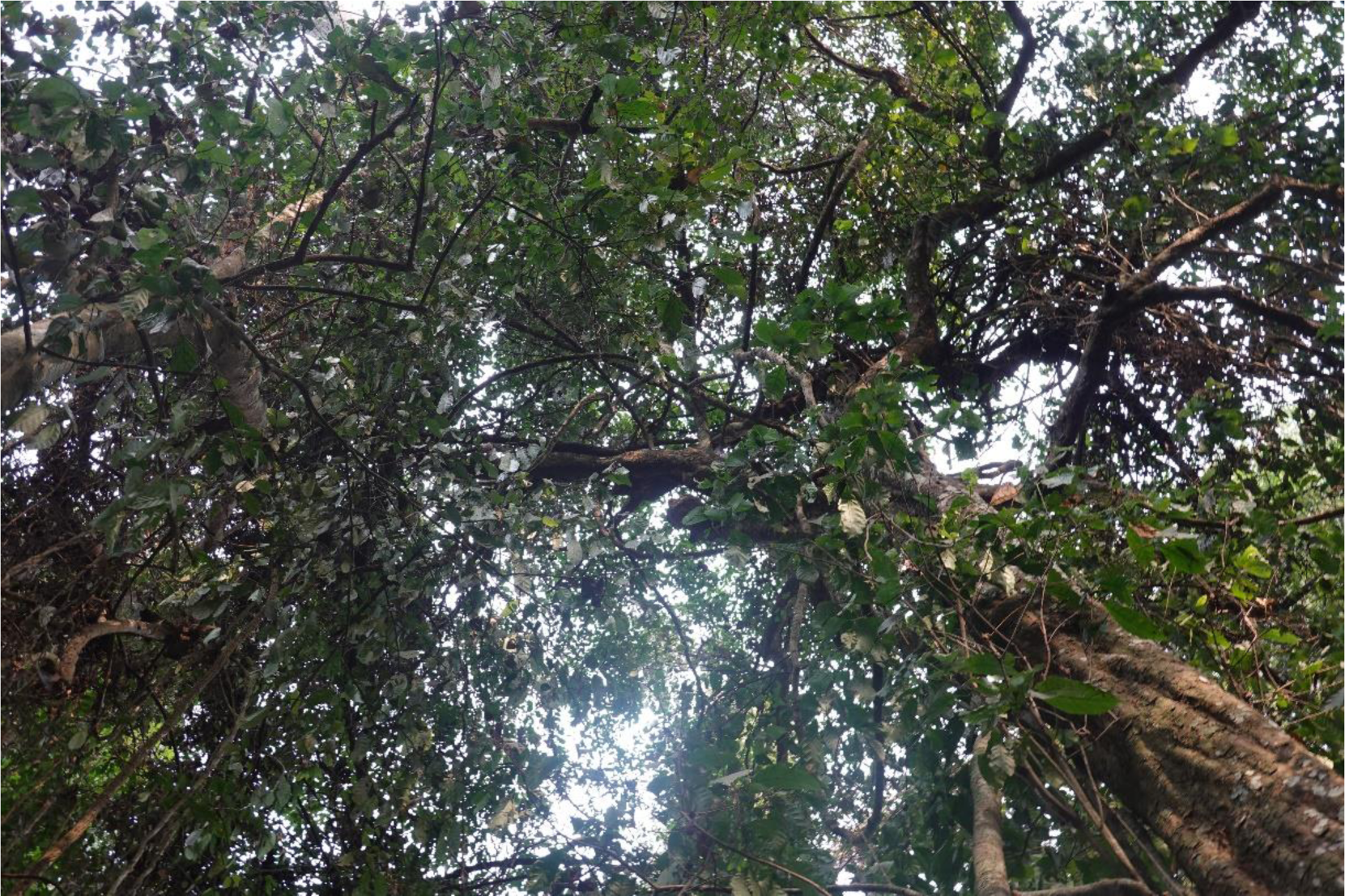

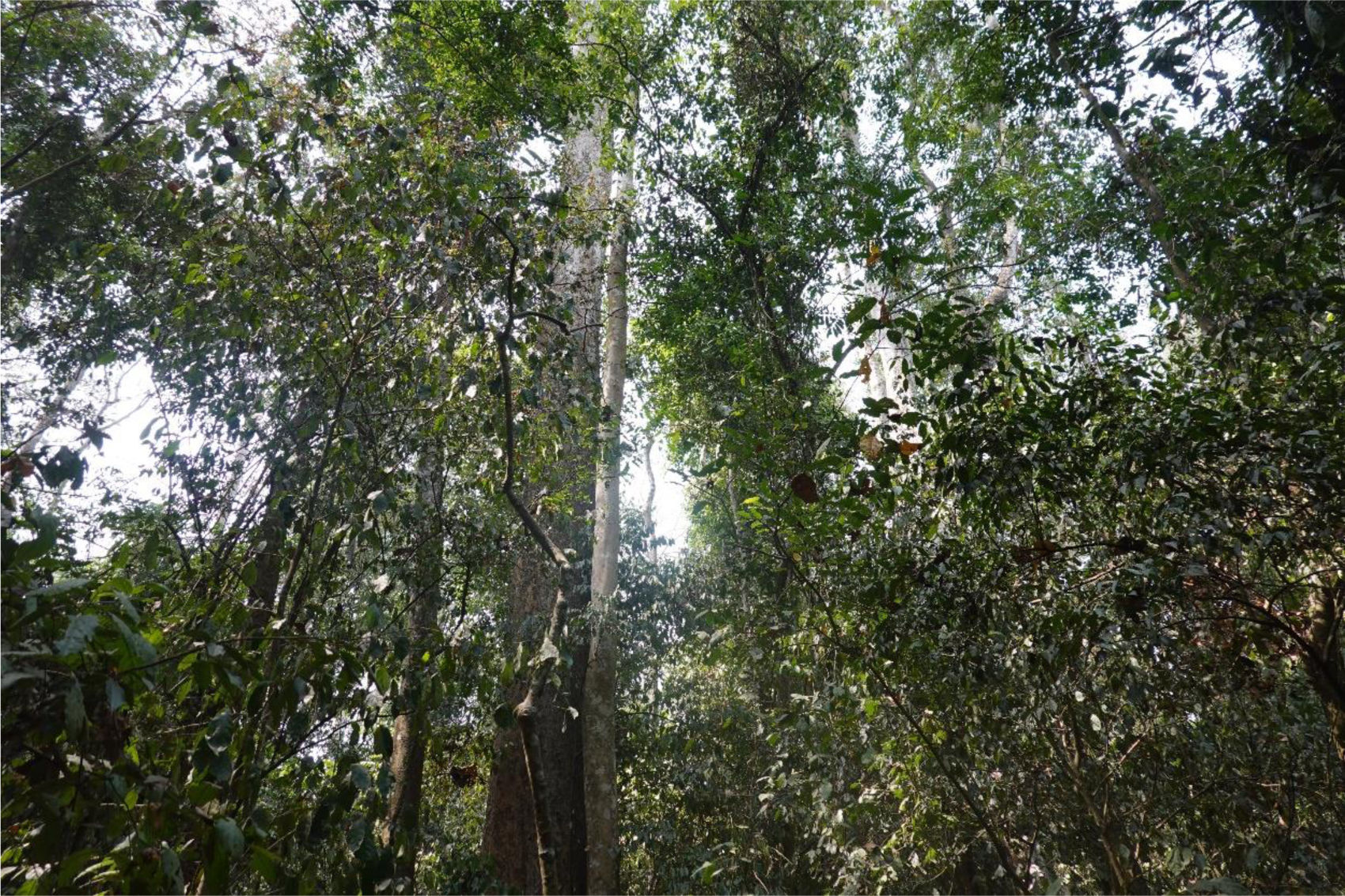

## Bobiri - BOB02

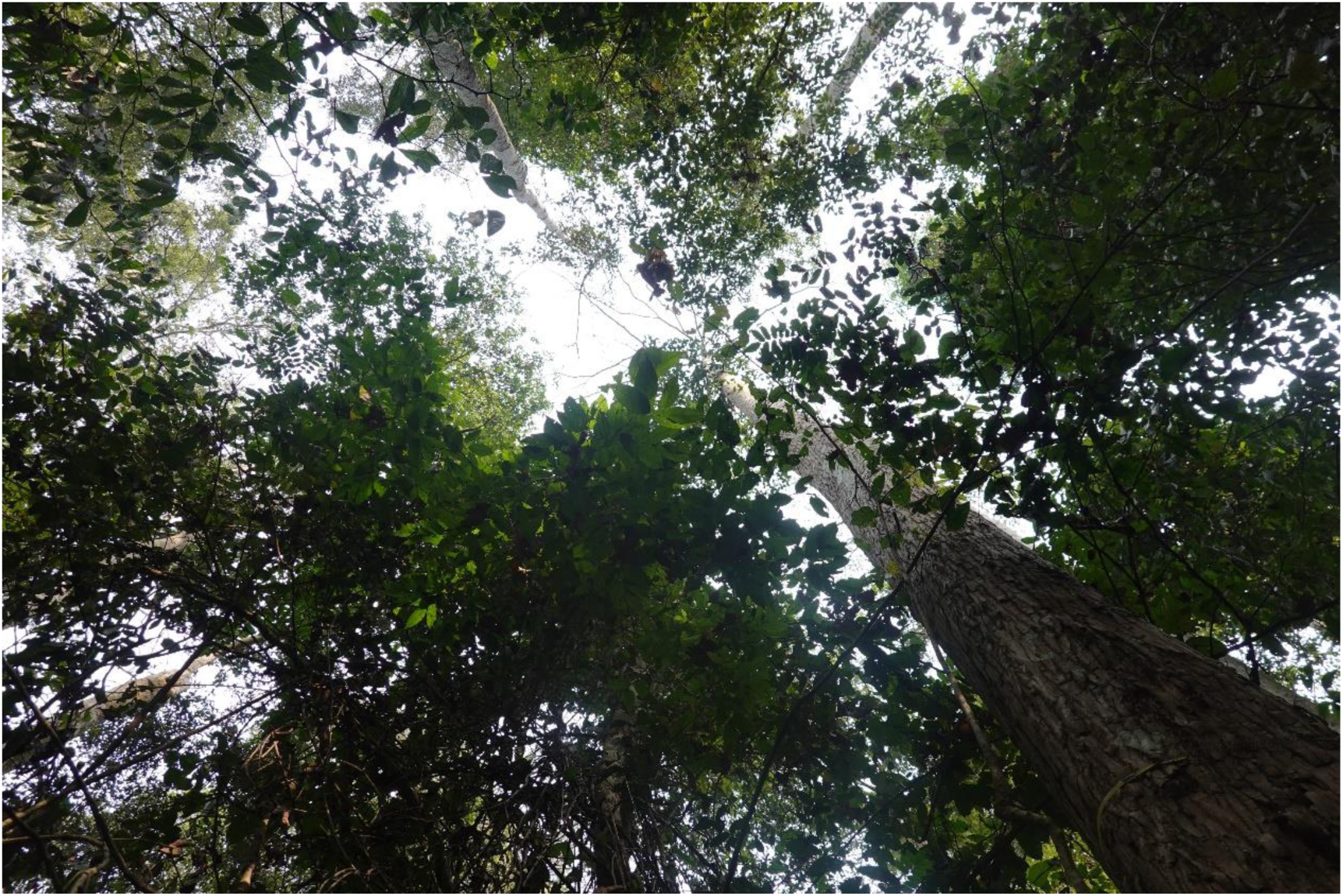

Photo Credit: all photos above were taken by Huanyuan Zhang-Zheng in January 2022.

## Kogaye - KOG01

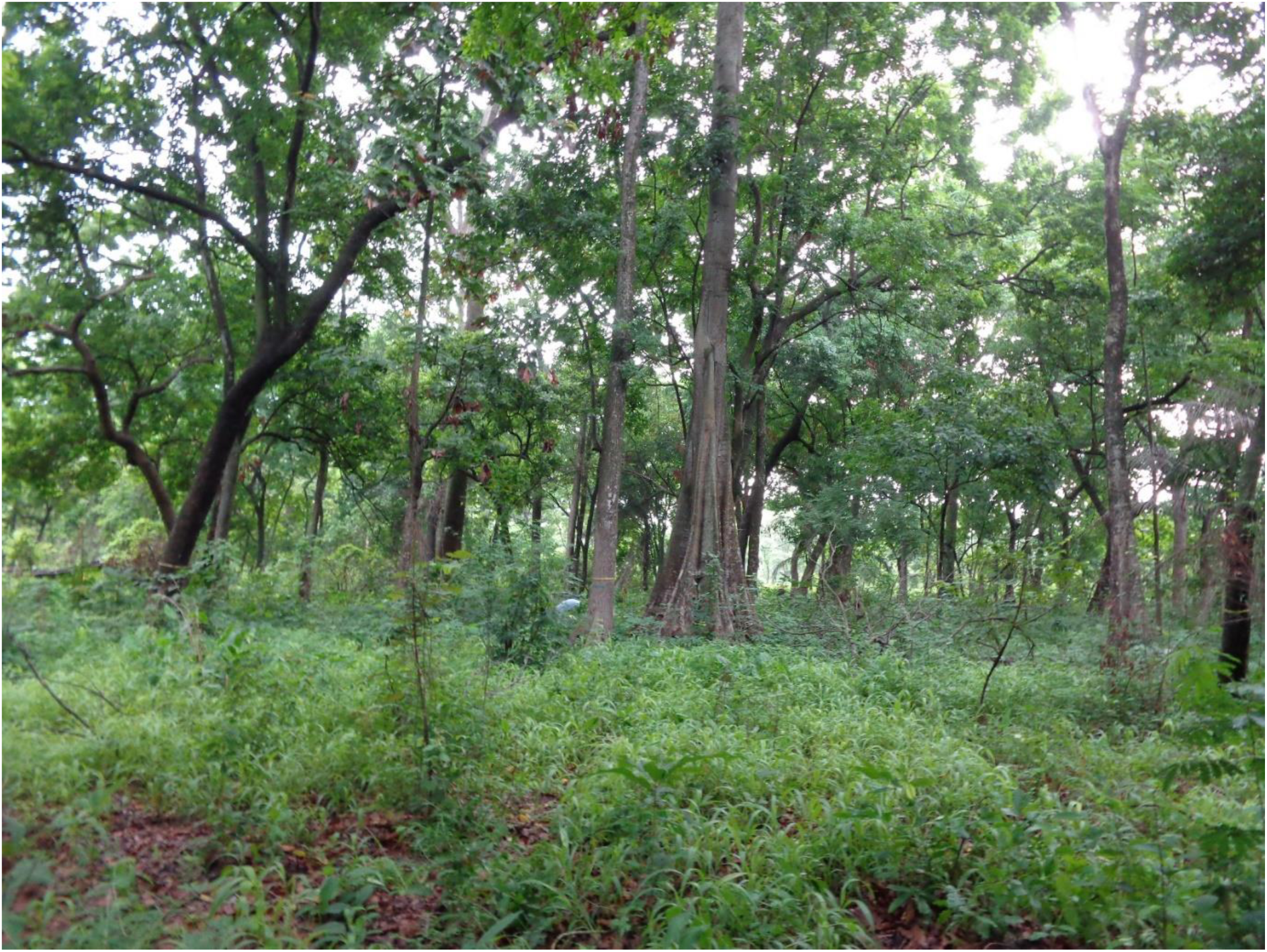

Photo Credit: the photo was shared by Akwasi Duah-Gyamfi. The photo was taken on 16 July 2013,

## Kogaye - KOG02

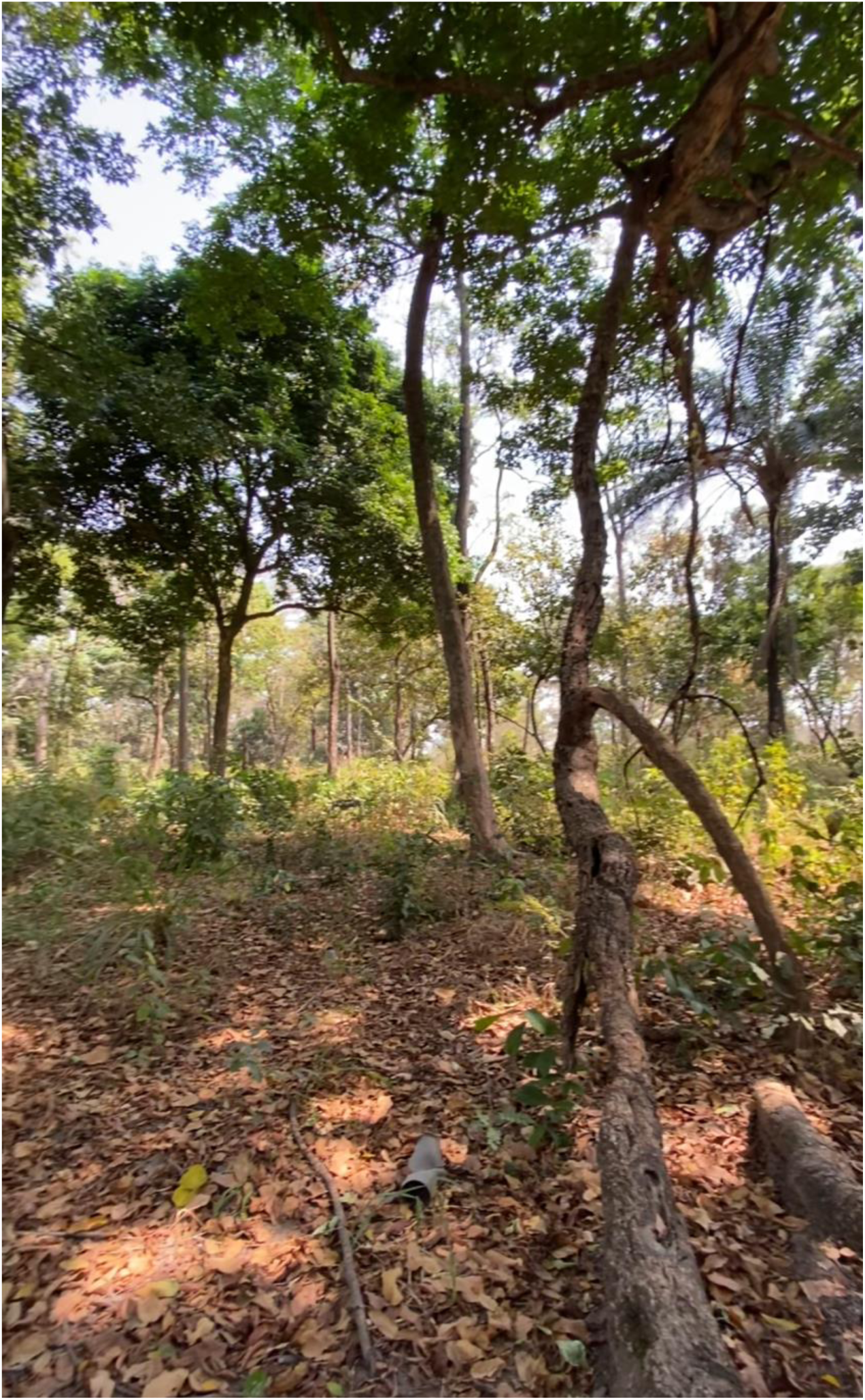

Photo Credit: taken by Huanyuan Zhang-Zheng in January 2022.

## Kogaye - KOG04

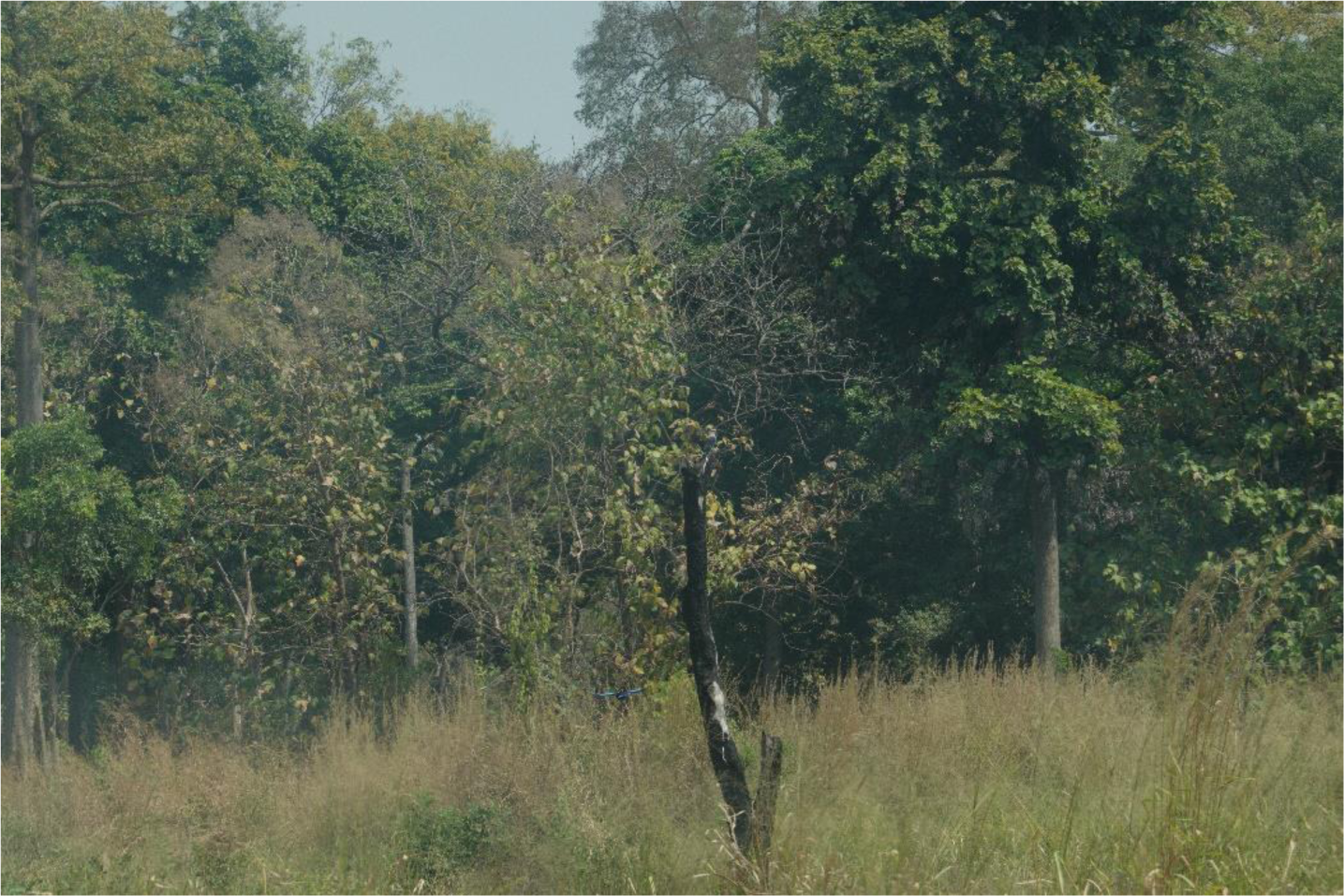

This is at the forest-savanna transition. Photo Credit: taken by Huanyuan Zhang-Zheng in January 2022.

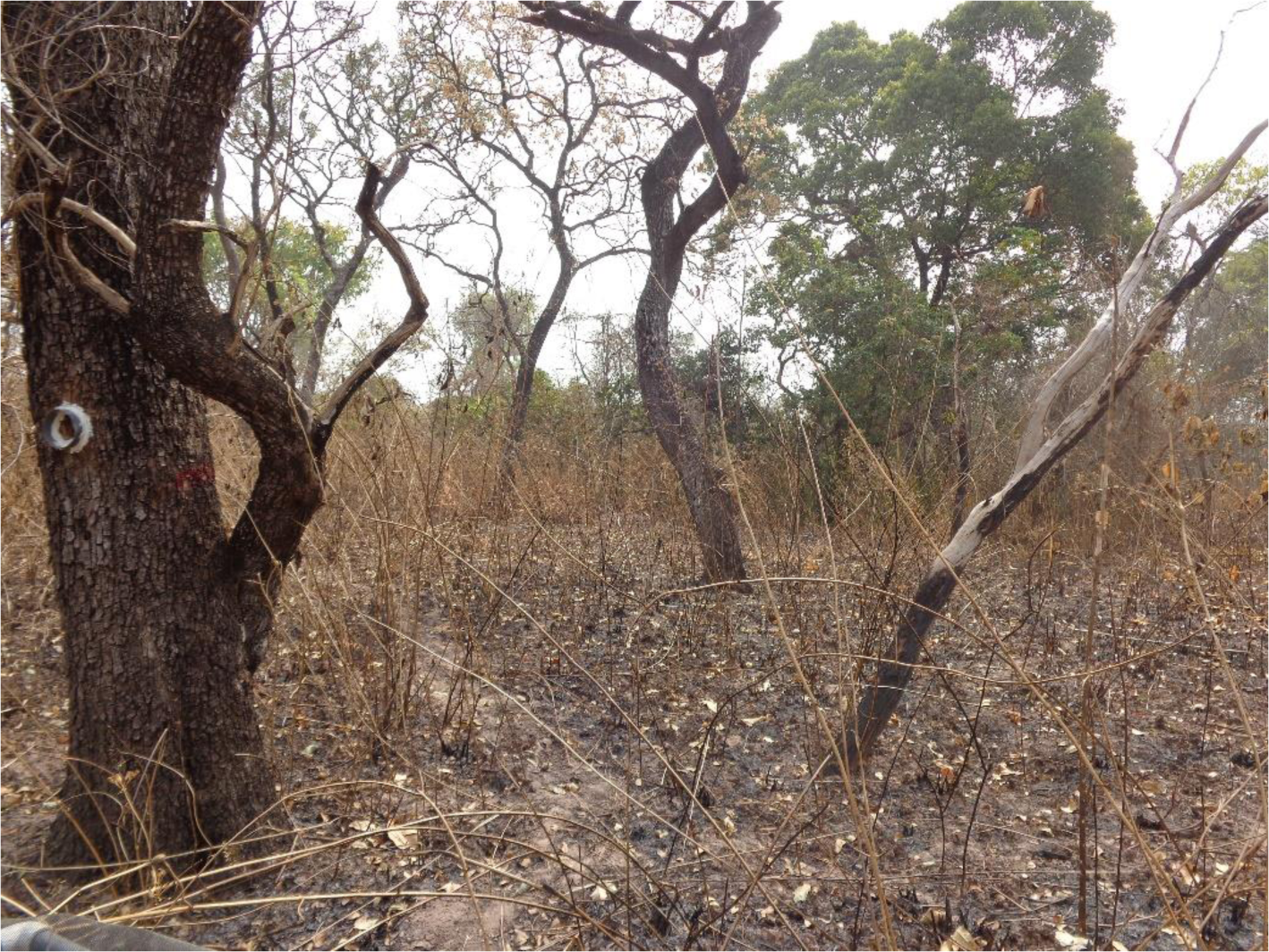

This plot rarely burns (as told by locals), but it looks like this when it does burn. Photo Credit: the photo was shared by Akwasi Duah-Gyamfi. The photo was taken on 03 February 2014,

## Kogaye - KOG05

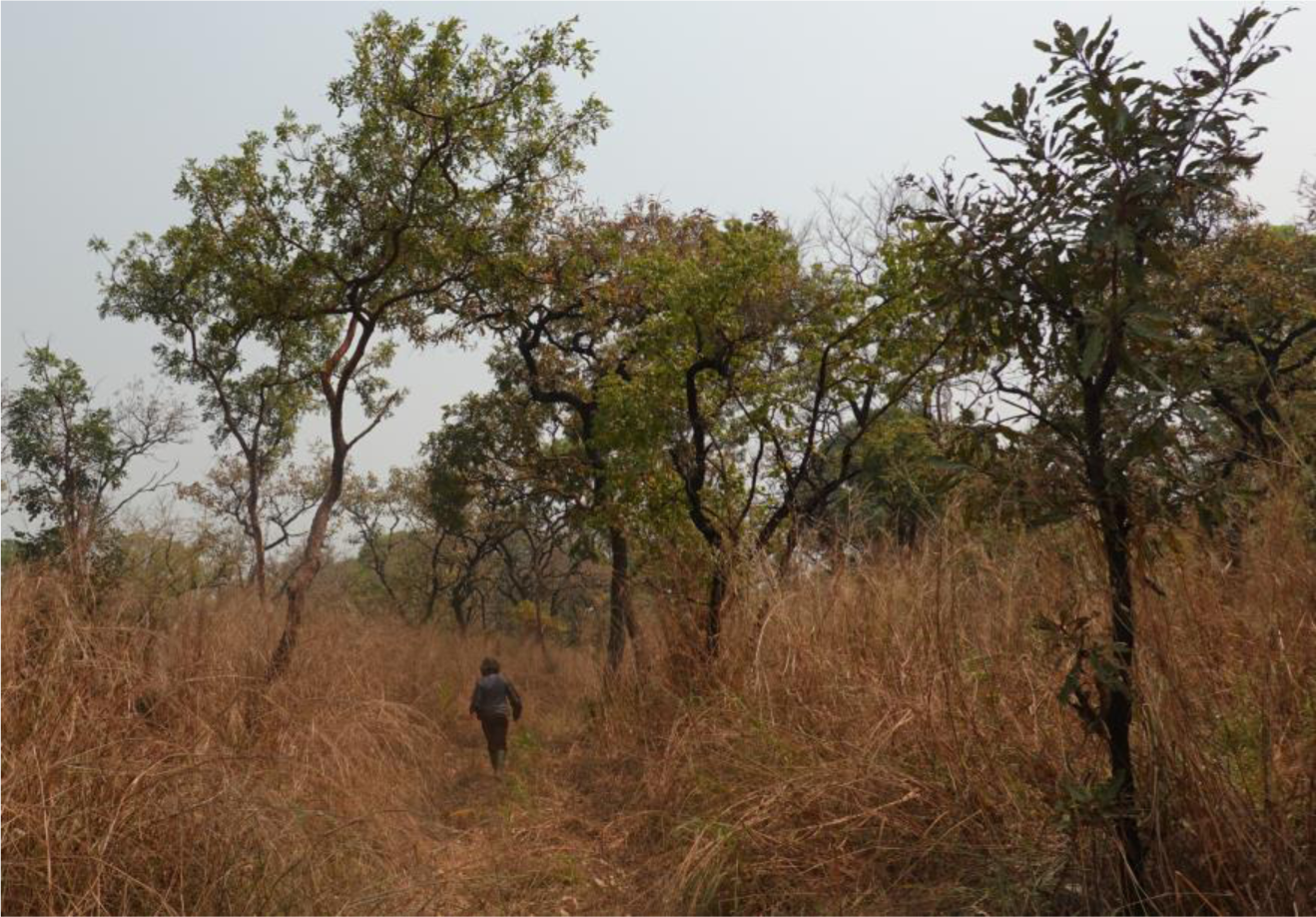

Photo Credit: taken by Huanyuan Zhang-Zheng in January 2022.

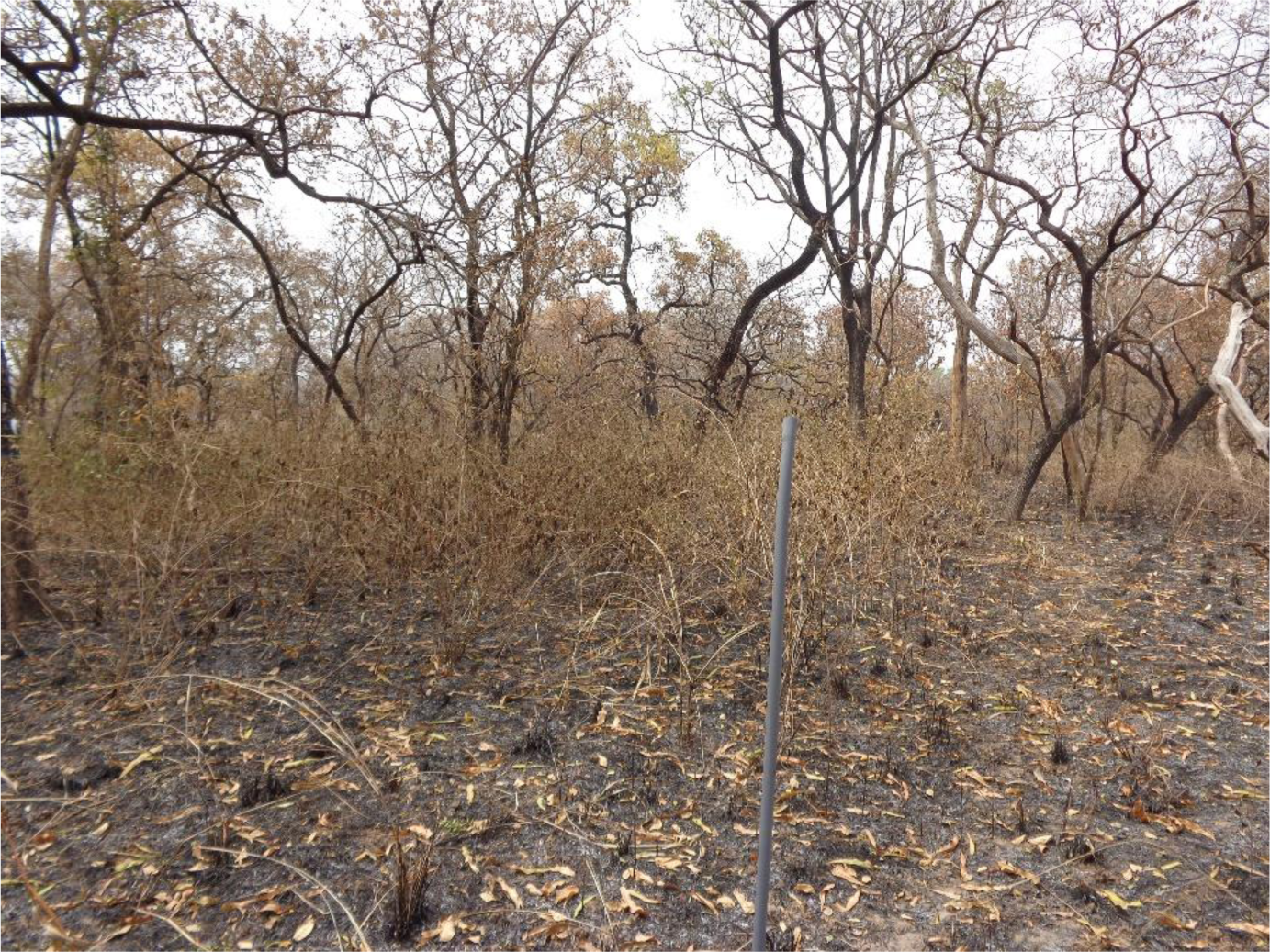

This plot frequently burns. Photo Credit: the photo was shared by Akwasi Duah-Gyamfi. The photo was taken on 06 February 2014,

## References

1. Grace, J., Malhi, Y., Meir, P. & Higuchi, N. Productivity of Tropical Rain Forests. In Terrestrial Global Productivity (2001). doi:10.1016/b978-012505290-0/50018-1.

2. Pan, Y. et al. A large and persistent carbon sink in the world’s forests. Science 333, (2011).

3. Liu, Y. Y. et al. Recent reversal in loss of global terrestrial biomass. Nat. Clim. Change 5, (2015).

4. Ciais, P. et al. Carbon and other biogeochemical cycles. in Climate change 2013: the physical science basis. Contribution of Working Group I to the Fifth Assessment Report of the Intergovernmental Panel on Climate Change 465–570 (Cambridge University Press, 2014).

5. Pugh, T. A. M. et al. A Large Committed Long-Term Sink of Carbon due to Vegetation Dynamics. Earths Future 6, (2018).

6. Jung, M. et al. Global patterns of land-atmosphere fluxes of carbon dioxide, latent heat, and sensible heat derived from eddy covariance, satellite, and meteorological observations. J. Geophys. Res. Biogeosciences 116, (2011).

7. Anav, A. et al. Spatiotemporal patterns of terrestrial gross primary production: A review. Rev. Geophys. 53, (2015).

8. Yang, R. et al. Divergent historical GPP trends among state-of-the-art multi-model simulations and satellite-based products. Earth Syst. Dyn. 13, (2022).

9. Bonan, G. B. et al. Improving canopy processes in the Community Land Model version 4 (CLM4) using global flux fields empirically inferred from FLUXNET data. J. Geophys. Res. 116, (2011).

10. Frankenberg, C. et al. New global observations of the terrestrial carbon cycle from GOSAT: Patterns of plant fluorescence with gross primary productivity. Geophys. Res. Lett. 38, (2011).

11. Chen, M. et al. Regional contribution to variability and trends of global gross primary productivity. Environ. Res. Lett. 12, 105005 (2017).

12. Badgley, G., Anderegg, L. D. L., Berry, J. A. & Field, C. B. Terrestrial gross primary production: Using NIRV to scale from site to globe. Glob. Change Biol. 25, (2019).

13. Hickler, T. et al. CO2 fertilization in temperate FACE experiments not representative of boreal and tropical forests. Glob. Change Biol. 14, (2008).

14. Wood, T. E., Cavaleri, M. A. & Reed, S. C. Tropical forest carbon balance in a warmer world: A critical review spanning microbial- to ecosystem-scale processes. Biol. Rev. 87, (2012).

15. Babst, F. et al. Modeling Ambitions Outpace Observations of Forest Carbon Allocation. Trends Plant Sci. 26, (2021).

16. Zhang, Y. & Ye, A. Would the obtainable gross primary productivity (GPP) products stand up? A critical assessment of 45 global GPP products. Sci. Total Environ. 783, 146965 (2021).

17. Joiner, J. et al. Estimation of terrestrial global gross primary production (GPP) with satellite data-driven models and eddy covariance flux data. Remote Sens. 10, 1346 (2018).

18. Tian, Z. et al. Fusion of multiple models for improving gross primary production estimation with eddy covariance data based on machine learning. J. Geophys. Res. Biogeosciences e2022JG007122 (2023).

19. Seiler, C., et al. Are terrestrial biosphere models fit for simulating the global land carbon sink? J. Adv. Model. Earth Syst. 14, e2021MS002946 (2022).

20. Ardö, J. Comparison between remote sensing and a dynamic vegetation model for estimating terrestrial primary production of Africa. Carbon Balance Manag. 10, (2015).

21. Zhang-Zheng, H. et al. Contrasting carbon cycle along tropical forest aridity gradients in W Africa and Amazonia. bioRxiv 2023–07 (2023).

22. Franklin, O. et al. Organizing principles for vegetation dynamics. Nat. Plants 6, 444–453 (2020).

23. Moore, S. et al. Forest biomass, productivity and carbon cycling along a rainfall gradient in West Africa. Glob. Change Biol. 24, e496–e510 (2018).

24. Gvozdevaite, A. et al. Leaf-level photosynthetic capacity dynamics in relation to soil and foliar nutrients along forest–savanna boundaries in Ghana and Brazil. Tree Physiol. 38, 1912–1925 (2018).

25. Oliveras, I. et al. The Influence of Taxonomy and Environment on Leaf Trait Variation Along Tropical Abiotic Gradients. Front. For. Glob. Change 3, 18 (2020).

26. Malhi, Y. et al. The Global Ecosystems Monitoring network: Monitoring ecosystem productivity and carbon cycling across the tropics. Biol. Conserv. 253, 108889 (2021).

27. Stocker, B. D. et al. P-model v1.0: An optimality-based light use efficiency model for simulating ecosystem gross primary production. Geosci. Model Dev. 13, 1545–1581 (2020).

28. Rogers, A. et al. A roadmap for improving the representation of photosynthesis in Earth system models. New Phytol. 213, 22–42 (2017).

29. Berg, S. et al. Ilastik: interactive machine learning for (bio) image analysis. Nat. Methods 16, 1226–1232 (2019).

30. Weiss, M. & Baret, F. CAN_EYE V6. 4.91 user manual. (2017).

31. Duursma, R. A. Plantecophys-an R package for analysing and modelling leaf gas exchange data. PloS One 10, e0143346 (2015).

32. De Kauwe, M. G. et al. A test of the ‘one-point method’for estimating maximum carboxylation capacity from field-measured, light-saturated photosynthesis. New Phytol. 210, 1130–1144 (2016).

33. Cornelissen, J. H. C. et al. A handbook of protocols for standardised and easy measurement of plant functional traits worldwide. Aust. J. Bot. 51, 335–380 (2003).

34. Keenan, T. F. & Niinemets, Ü. Global leaf trait estimates biased due to plasticity in the shade. Nat. Plants 3, 1–6 (2016).

35. Friedlingstein, P. et al. Global Carbon Budget 2020. Earth Syst. Sci. Data 12, (2020).

36. Jung, M. et al. The FLUXCOM ensemble of global land-atmosphere energy fluxes. Sci. Data 6, 74 (2019).

37. Running, S., Mu, Q. & Zhao, M. MOD17A2H MODIS/Terra Gross Primary Productivity 8-Day L4 Global 500m SIN Grid V006. NASA EOSDIS Land Processes DAAC 10.5067/MODIS/MOD17A2H.006 (2015).

38. Farquhar, G. D., von Caemmerer, S. & Berry, J. A. A biochemical model of photosynthetic CO2 assimilation in leaves of C3 species. Planta 149, 78–90 (1980).

39. Hilker, T., Coops, N. C., Wulder, M. A., Black, T. A. & Guy, R. D. The use of remote sensing in light use efficiency based models of gross primary production: A review of current status and future requirements. Sci. Total Environ. 404, 411–423 (2008).

40. Sun, Y. et al. Impact of mesophyll diffusion on estimated global land CO2 fertilization. Proc. Natl. Acad. Sci. 111, 15774–15779 (2014).

41. Harrison, S. P. et al. Eco-evolutionary optimality as a means to improve vegetation and land-surface models. New Phytol. 231, 2125–2141 (2021).

42. Friend, A. Modelling canopy CO2 fluxes: are ‘big-leaf’simplifications justified? Glob. Ecol. Biogeogr. 10, 603–619 (2001).

43. Wang, H. et al. Towards a universal model for carbon dioxide uptake by plants. Nat. Plants 2017 39 3, 734–741 (2017).

44. Smith, N. G. et al. Global photosynthetic capacity is optimized to the environment. (2019) doi:10.1111/ele.13210.

45. Peng, Y., Bloomfield, K. J., Cernusak, L. A., Domingues, T. F. & Colin Prentice, I. Global climate and nutrient controls of photosynthetic capacity. *Commun*. Biol. 2021 41 4, 1–9 (2021).

46. Peng, Y., Bloomfield, K. J. & Prentice, I. C. A theory of plant function helps to explain leaf-trait and productivity responses to elevation. New Phytol. 226, 1274–1284 (2020).

47. Morel, A. C. et al. Carbon dynamics, net primary productivity and human-appropriated net primary productivity across a forest–cocoa farm landscape in West Africa. Glob. Change Biol. 25, (2019).

48. Luo, X. et al. Comparison of Big-Leaf, Two-Big-Leaf, and Two-Leaf Upscaling Schemes for Evapotranspiration Estimation Using Coupled Carbon-Water Modeling. J. Geophys. Res. Biogeosciences 123, 207–225 (2018).

49. Domingues, T. F. et al. Co-limitation of photosynthetic capacity by nitrogen and phosphorus in West Africa woodlands. Plant Cell Environ. 33, 959–980 (2010).

50. Tomlinson, K. W. et al. Leaf adaptations of evergreen and deciduous trees of semi-arid and humid savannas on three continents. J. Ecol. 101, 430–440 (2013).

51. Xu, L. et al. Satellite observation of tropical forest seasonality: spatial patterns of carbon exchange in Amazonia. Environ. Res. Lett. 10, 084005 (2015).

52. Rödig, E. et al. The importance of forest structure for carbon fluxes of the Amazon rainforest. Environ. Res. Lett. 13, 054013 (2018).

53. Zhao, M., Heinsch, F. A., Nemani, R. R. & Running, S. W. Improvements of the MODIS terrestrial gross and net primary production global data set. Remote Sens. Environ. 95, (2005).

54. Fensholt, R. & Proud, S. R. Evaluation of Earth Observation based global long term vegetation trends - Comparing GIMMS and MODIS global NDVI time series. Remote Sens. Environ. 119, 131–147 (2012).

55. Fuster, B. et al. Quality Assessment of PROBA-V LAI, fAPAR and fCOVER Collection 300 m Products of Copernicus Global Land Service. Remote Sens. 12, 1017 (2020).

56. Sudmanns, M., Tiede, D., Augustin, H. & Lang, S. Assessing global Sentinel-2 coverage dynamics and data availability for operational Earth observation (EO) applications using the EO-Compass. Int. J. Digit. Earth 13, 768–784 (2020).

57. Wang, L. et al. Evaluation of the latest MODIS GPP products across multiple biomes using global eddy covariance flux data. Remote Sens. 9, (2017).

58. Chen, J. M. et al. Global datasets of leaf photosynthetic capacity for ecological and earth system research. Earth Syst. Sci. Data 14, 4077–4093 (2022).

59. Dong, N. et al. Rising CO2 and warming reduce global canopy demand for nitrogen. New Phytol. 235, 1692–1700 (2022).

60. Sibret, T. et al. High photosynthetic capacity of Sahelian C3 and C4 plants. Photosynth. Res. 147, 161–175 (2021).

61. Lu, X., Croft, H., Chen, J. M., Luo, Y. & Ju, W. Estimating photosynthetic capacity from optimized Rubisco–chlorophyll relationships among vegetation types and under global change. Environ. Res. Lett. 17, 014028 (2022).

62. Rogers, A. The use and misuse of Vc,max in Earth System Models. Photosynth. Res. 119, 15–29 (2014).

63. Kattge, J., Knorr, W., Raddatz, T. & Wirth, C. Quantifying photosynthetic capacity and its relationship to leaf nitrogen content for global-scale terrestrial biosphere models. Glob. Change Biol. 15, 976–991 (2009).

64. Belelli Marchesini, L., et al. Ankasa Flux Tower: A New Research Facility for the Study of the Carbon Cycle in a Primary Tropical Forest in Africa. (2011).

65. Pastorello, G. et al. The FLUXNET2015 dataset and the ONEFlux processing pipeline for eddy covariance data. Sci. Data 7, 225 (2020).

66. Hayek, M. N. et al. A novel correction for biases in forest eddy covariance carbon balance. Agric. For. Meteorol. 250–251, 90–101 (2018).

67. Billesbach, D. P. et al. Effects of the Gill-Solent WindMaster-Pro “w-boost” firmware bug on eddy covariance fluxes and some simple recovery strategies. Agric. For. Meteorol. 265, 145–151 (2019).

68. de Araújo, A. C. et al. The spatial variability of CO2 storage and the interpretation of eddy covariance fluxes in central Amazonia. Agric. For. Meteorol. 150, 226–237 (2010).

69. Fu, Z. et al. The surface-atmosphere exchange of carbon dioxide in tropical rainforests: Sensitivity to environmental drivers and flux measurement methodology. Agric. For. Meteorol. 263, 292–307 (2018).

70. Aguirre-Gutiérrez, J. et al. Pantropical modelling of canopy functional traits using Sentinel-2 remote sensing data. Remote Sens. Environ. 252, 112122 (2021).

71. Wullschleger, S. D. et al. Plant functional types in Earth system models: past experiences and future directions for application of dynamic vegetation models in high-latitude ecosystems. Ann. Bot. 114, 1–16 (2014).

72. Prentice, I. C., Liang, X., Medlyn, B. E. & Wang, Y. P. Reliable, robust and realistic: The three R’s of next-generation land-surface modelling. Atmospheric Chem. Phys. 15, 5987–6005 (2015).

73. Buchanan, G. M., Field, R. H., Bradbury, R. B., Luraschi, B. & Vickery, J. A. The impact of tree loss on carbon management in West Africa. Carbon Manag. 12, 623–633 (2021).

74. Hungate, B. A., Dukes, J. S., Shaw, M. R., Luo, Y. & Field, C. B. Nitrogen and Climate Change. Science 302, (2003).

75. Zhang-Zheng, H. et al. Photosynthetic and water transport strategies of plants along a tropical forest aridity gradient: a test of optimality theory. 2023.01.10.523419 Preprint at 10.1101/2023.01.10.523419 (2023).

76. Chen, Y., Feng, X., Fu, B., Wu, X. & Gao, Z. Improved Global Maps of the Optimum Growth Temperature, Maximum Light Use Efficiency, and Gross Primary Production for Vegetation. J. Geophys. Res. Biogeosciences 126, e2020JG005651 (2021).

77. Cornwell, W. K. et al. A global dataset of leaf delta 13C values. Sci. Data (2016).

78. Burton, C., Rifai, S. & Malhi, Y. Inter-comparison and assessment of gridded climate products over tropical forests during the 2015/2016 El Niño. Philos. Trans. R. Soc. B Biol. Sci. 373, 20170406 (2018).

79. Jin, W. et al. Leaf development and demography explain photosynthetic seasonality in Amazon evergreen forests. Science 351, 972–976 (2016).

80. Zhang, Y., Chen, J. M. & Miller, J. R. Determining digital hemispherical photograph exposure for leaf area index estimation. Agric. For. Meteorol. 133, 166–181 (2005).

81. Zhang, Y. et al. Modeling the impacts of diffuse light fraction on photosynthesis in ORCHIDEE (v5453) land surface model. Geosci. Model Dev. 13, 5401–5423 (2020).

82. Madansky, A. & Alexander, H. Weighted standard error and its impact on significance testing. Anal. Group Inc (2017).

83. Bernacchi, C., Pimentel, C. & Long, S. P. In vivo temperature response functions of parameters required to model RuBP-limited photosynthesis. Plant Cell Environ. 26, 1419–1430 (2003).

84. Wang, H. et al. Photosynthetic responses to altitude: an explanation based on optimality principles. New Phytol. 213, 976–982 (2017).

85. Kou-Giesbrecht, S. & Arora, V. K. Representing the Dynamic Response of Vegetation to Nitrogen Limitation via Biological Nitrogen Fixation in the CLASSIC Land Model. Glob. Biogeochem. Cycles 36, e2022GB007341 (2022).

